## Supplementary Material for "Molecular Architecture of Chitin and Chitosan-Dominated Cell Walls in Zygomycetous Fungal Pathogens by Solid-State NMR"

### Table of Contents


Supplementary Fig. 1. Replication of *R. delemar* samples 3
Supplementary Fig. 2. Resonance assignment of chitin, chitosan, and glucan in *R. delemar* 4
Supplementary Fig. 3. Rigid cell wall polysaccharides in *R. delemar* 5
Supplementary Fig. 4. Biopolymer signals resolved in 2D $^{13}$C refocused J-INADEQUATE 6
Supplementary Fig. 5. Distribution of chitin, chitosan, and β-glucan in rigid and mobile domains 7
Supplementary Fig. 6. Identification of mobile nitrogenated carbohydrates 8
Supplementary Fig. 7. Carbon connectivity of five types of fucose residues 9
Supplementary Fig. 8. Absence of galacturonic acid and glucuronic acid in *R. delemar* 10
Supplementary Fig. 9. β-glucans in the mobile domain of nikkomycin-treated *R. delemar* cell wall 11
Supplementary Fig. 10. Chitin is the only carbohydrate detectable in 15 ms PAR spectrum 12
Supplementary Fig. 11. Intermolecular contacts observed in nikkomycin-treated *R. delemar* 13
Supplementary Fig. 12. Proteins are partially ordered and closely packed in *R. delemar*, apo 14
Supplementary Fig. 13. Presence of protein in both mobile and rigid domains 15
Supplementary Fig. 14. Water-edited experiment setup for inspecting carbohydrate hydration 16
Supplementary Fig. 15. Nikkomycin alters water accessibility of rigid polysaccharides 17
Supplementary Fig. 16. $^{13}$C-$T_1$ relaxation in apo and nikkomycin-treated *R. delemar* 18
Supplementary Fig. 17. $^{1}$H-$T_{1\rho}$ relaxation in *R. delemar* 19
Supplementary Fig. 18. Distinct spectral patterns of *Rhizopus*, *Mucor*, and *Aspergillus* 20
Supplementary Fig. 19. Similarity of *Rhizopus* and *Mucor* carbohydrates 21
Supplementary Fig. 20. Experimental flowchart of fungal cultivation 22
Supplementary Fig. 21. Experimental flowchart of constructing fungal growth curves 23
Supplementary Table 1. Measurements of the cell-wall thickness of *R. delemar* 24
Supplementary Table 2. Solid-state NMR experimental parameters 25
Supplementary Table 3. $^{13}$C and $^{15}$N chemical shifts of rigid polysaccharides in *R. delemar* 26
Supplementary Table 4. $^{13}$C chemical shifts of mobile polysaccharides in *R. delemar* 27
Supplementary Table 5. Chemical shifts of amino acids in the fungal cells 28
Supplementary Table 6. Molar composition of polysaccharides in cell walls 29
Supplementary Table 7. Intermolecular cross peaks probed by ssNMR 30
Supplementary Table 8. Water-edited intensities of polysaccharide carbon sites 31
Supplementary Table 9. $^{13}$C-$T_1$ and $^{1}$H-$T_{1\rho}$ relaxation time constants 32
Supplementary References 33

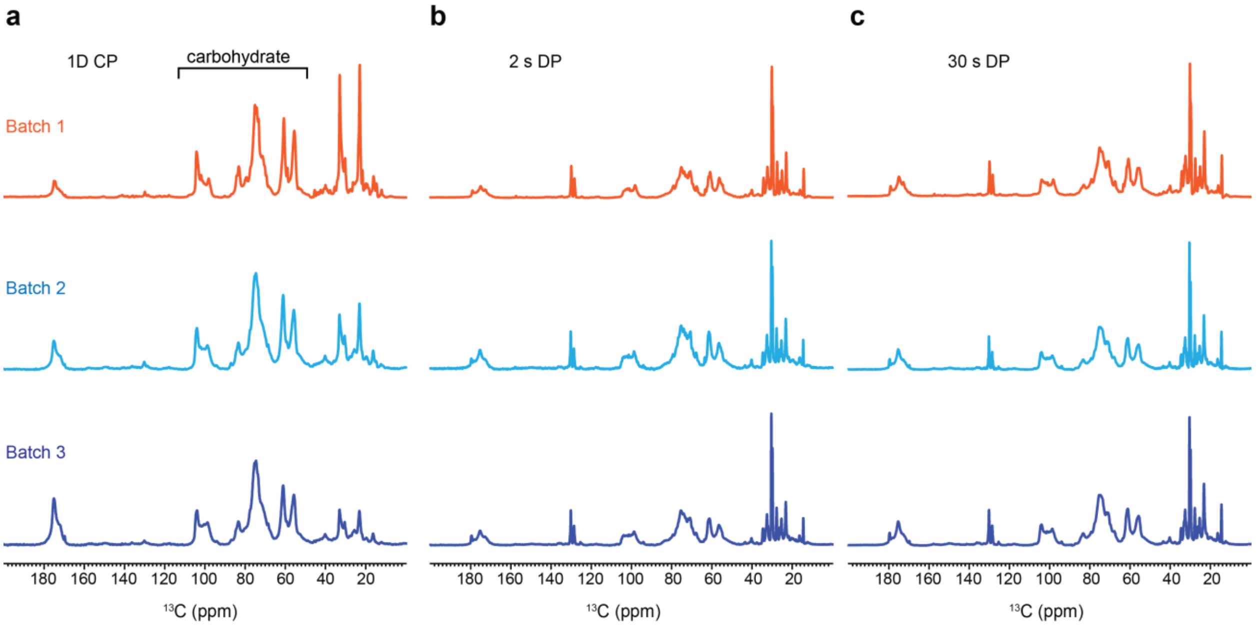


**Supplementary Figure 1. Replication of *R. delemar* samples.** (**a**) 1D $^{13}$C CP spectra of apo *R. delemar*. (**b**) Mobile components detected by 1D $^{13}$C DP with a short recycle delay of 2 s. (**c**) Quantitative $^{13}$C DP spectra for the detection of all molecules through a recycle delay of 30 s. The spectra exhibit high reproducibility across three separate batches prepared in 2018 (batch 1), 2022 (batch 2) and 2023 (batch 3).

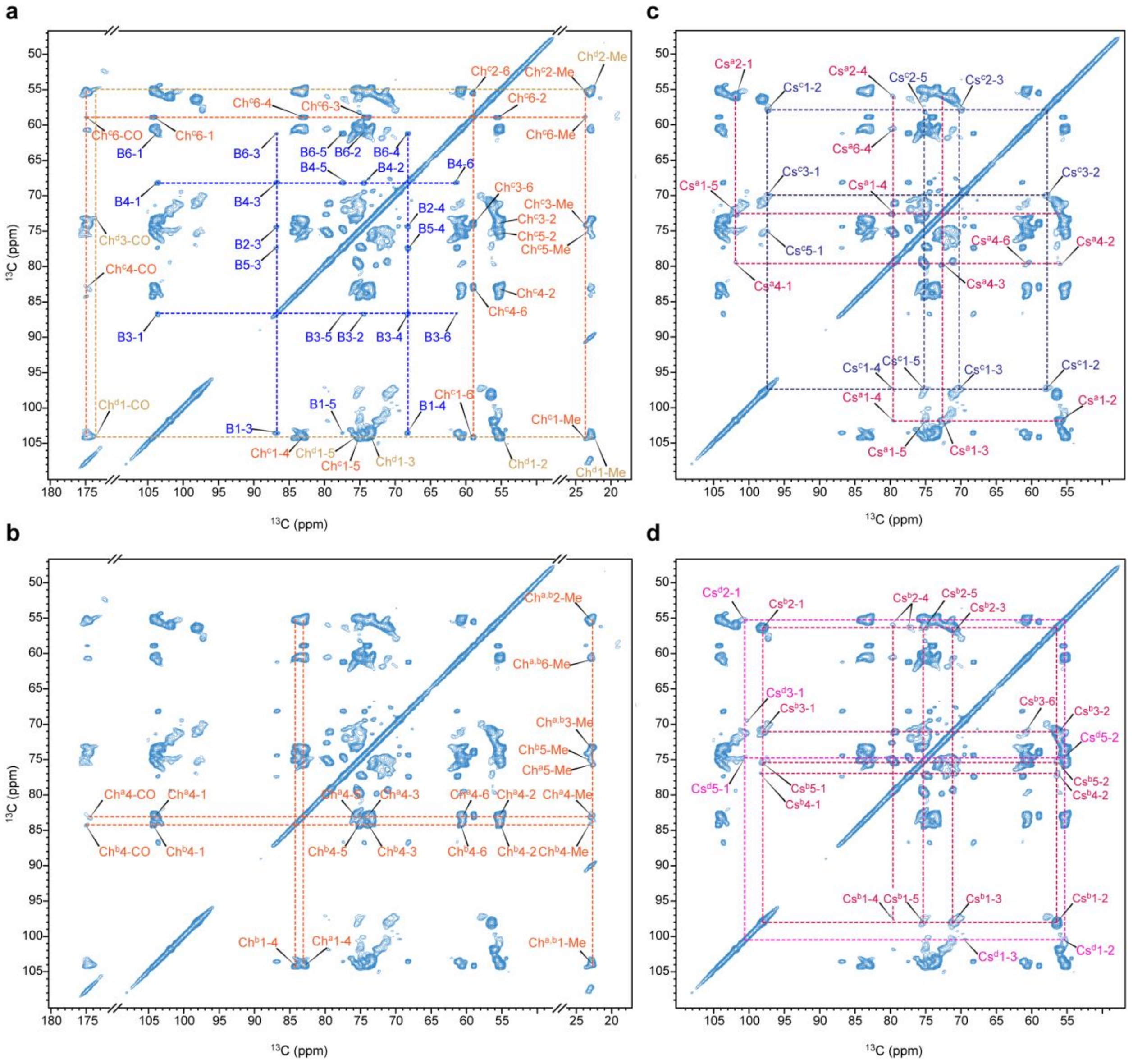


**Supplementary Figure 2. Resonance assignment of chitin, chitosan, and glucan in *R. delemar* cell wall.** CP-based 2D $^{13}$C-$^{13}$C correlation spectrum measured with 53 ms CORD mixing were plotted as four separate panels to show the well-resolved signals of **(a)** β-1,3-glucan and type-c and type-d of chitin forms, **(b)** type-a and type-b chitin, **(c)** type-a and type-c chitosan molecules, and **(d)** type-b and type-d chitosan molecules. NMR abbreviations are used for β-1,3-glucan (B), chitin (Ch), and chitosan (Cs). Superscripts indicate the subform type within each molecule. Carbon numbers are also labeled for each cross peak. For example, Ch$^c$2-Me indicates the cross peak between carbon 2 and methyl carbon within type-c chitin.

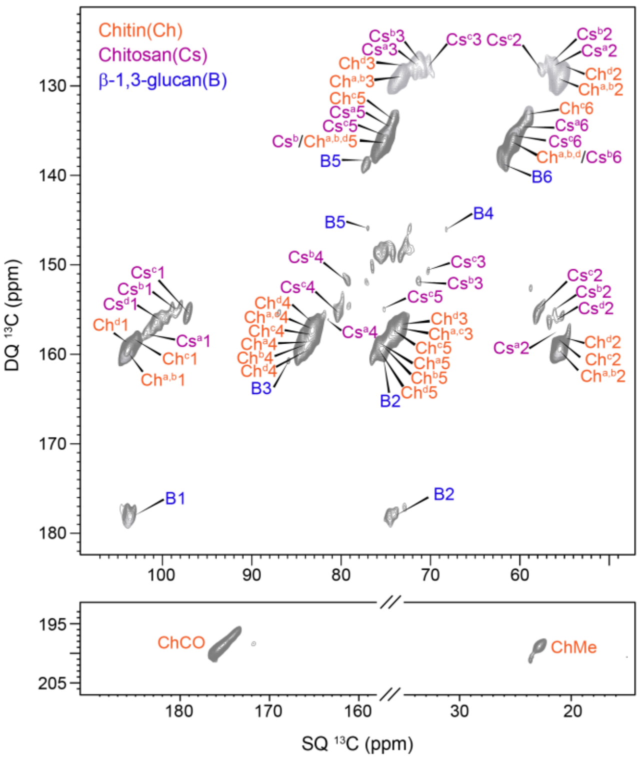


**Supplementary Figure 3. Rigid cell wall polysaccharides in *R. delemar*.** Through-bond carbon connectivity was resolved using 2D $^{13}$C CP refocused J-INADEQUATE spectrum. Multiple sets of conformers were resolved for chitin and chitosan, in addition to a single type of β-1,3-glucan. Each peak is annotated with the abbreviation of the carbohydrate name, the subtype (in superscript), and the carbon number. For instance, Ch$^{c}$5 represents the carbon 5 of type-c chitin. The linkage between the carbonyl (CO) and methyl (Me) of chitin is shown in the bottom panel.

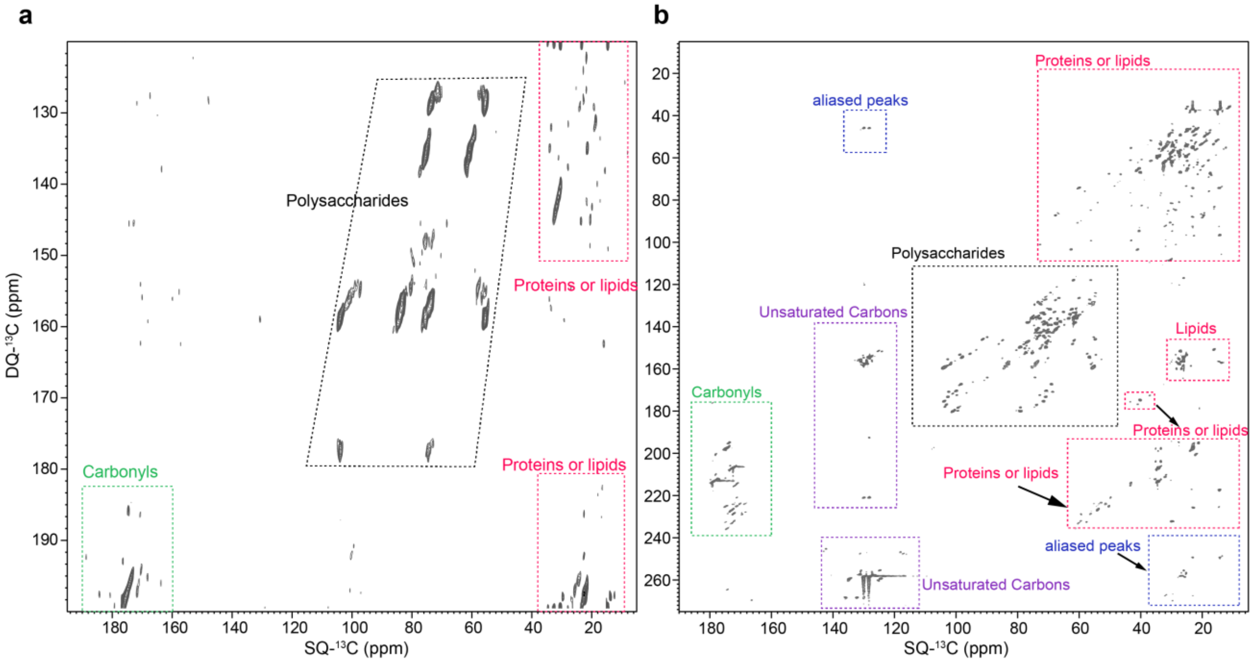


**Supplementary Figure 4. Biopolymer signals resolved in 2D $^{13}$C refocused J-INADEQUATE**. (**a**) Rigid components of the cell wall in apo *R. delemar* sample probed using 2D CP $^{13}$C refocused J-INADEQUAT E spectrum. The individual regions were marked with different colors of quadrangles: green for carbonyls, black for polysaccharides, and red for aliphatics from proteins and lipids. (**b)** Mobile components of apo *R. delemar* sample detected using 2D DP $^{13}$C refocused J-INADEQUATE spectrum. The individual regions are marked using the same color code mentioned above. The unsaturated carbons (aromatics or C=C) are marked in purple and the aliased peaks are boxed in blue.

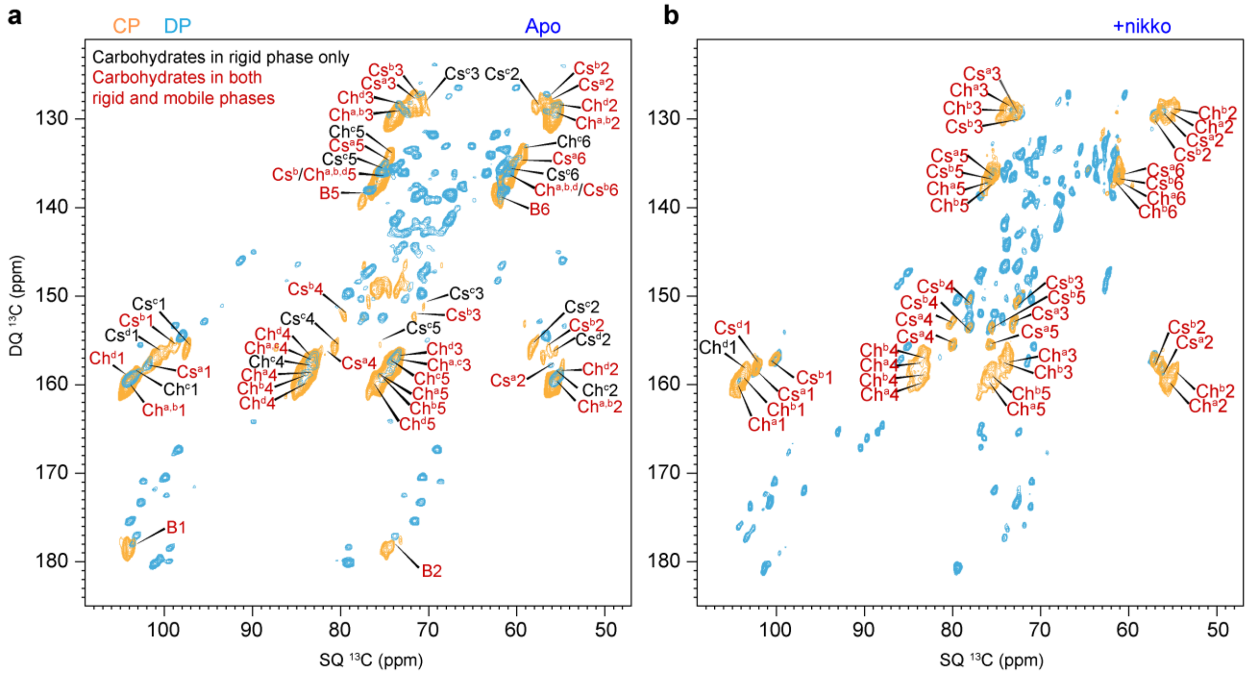


**Supplementary Figure 5. Distribution of chitin, chitosan, and β-glucan in rigid and mobile domains**. Overlay of 2D refocused J-INADEQUATE spectra measured with CP (orange) and DP (cyan) for (**a**) apo and (**b**) nikkomycin-treated *R. delemar* samples. The carbohydrates observed only in the CP-based spectra are rigid and are marked in black. The carbohydrates observed in both CP (INADEQUATE here and CORD in Figure S2) and DP-based spectra have two-modal distribution in rigid and mo bile phases, and are marked in red. Types-a,b chitin, types-a,b chitosan and β-1,3-glucan were observed in both domains for both samples. The Peak (DQ~159.8 ppm, SQ~104.1 ppm) of the $^{13}$C DP J-INADEQUATE spectra (blue) in panel b are assigned as types-a,b chitin, but other carbons of them (C3, C4, C5, and C6) are not confirmed.

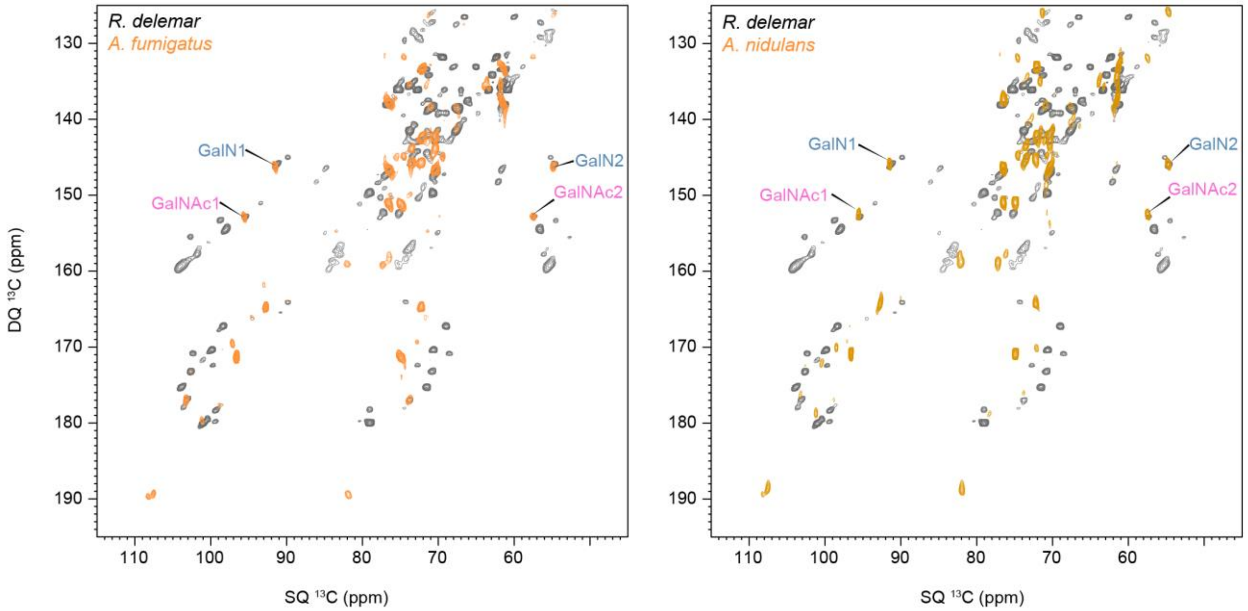


**Supplementary Figure 6. Identification of mobile nitrogenated carbohydrates**. One type of GalNAc and GalN are respectively confirmed by overlapping the 2D DP $^{13}$C J-INADEQUATE spectrum of *R. delemar* with two *Aspergillus* species (*A. fumigatus* and *A. nidulans*). The signals of galactosamine (GalN) and N-acetylgalactosamine (GalNAc) present in the mobile region of *R. delemar* are well-matched with the two A*spergillus* samples.

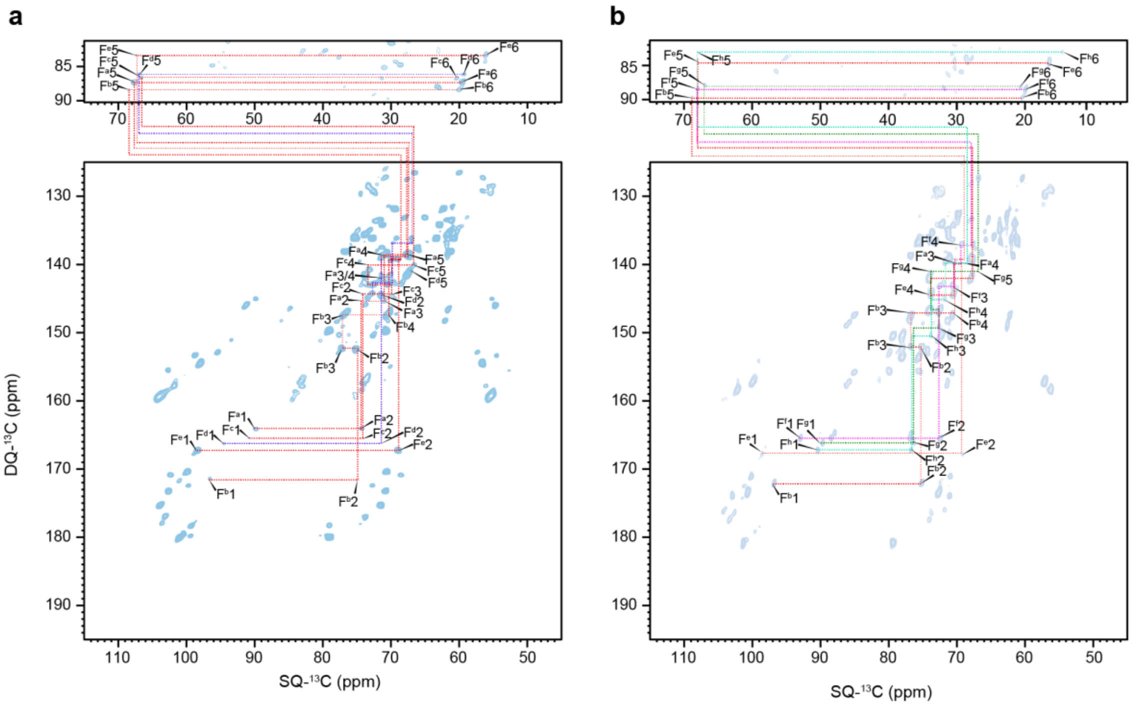


**Supplementary Figure 7. Carbon connectivity of five types of fucose residues.** The connectivity is tracked starting from the unique C5-C6 pair that involves a methyl carbon and a carbon on the pyranose ring of the carbohydrate. Similar types of fucose residues can be found in the $^{13}$C DP refocused INADEQUATE spectra of both (**a**) apo and (**b**) nikkomycin-treated *R. delemar* samples.

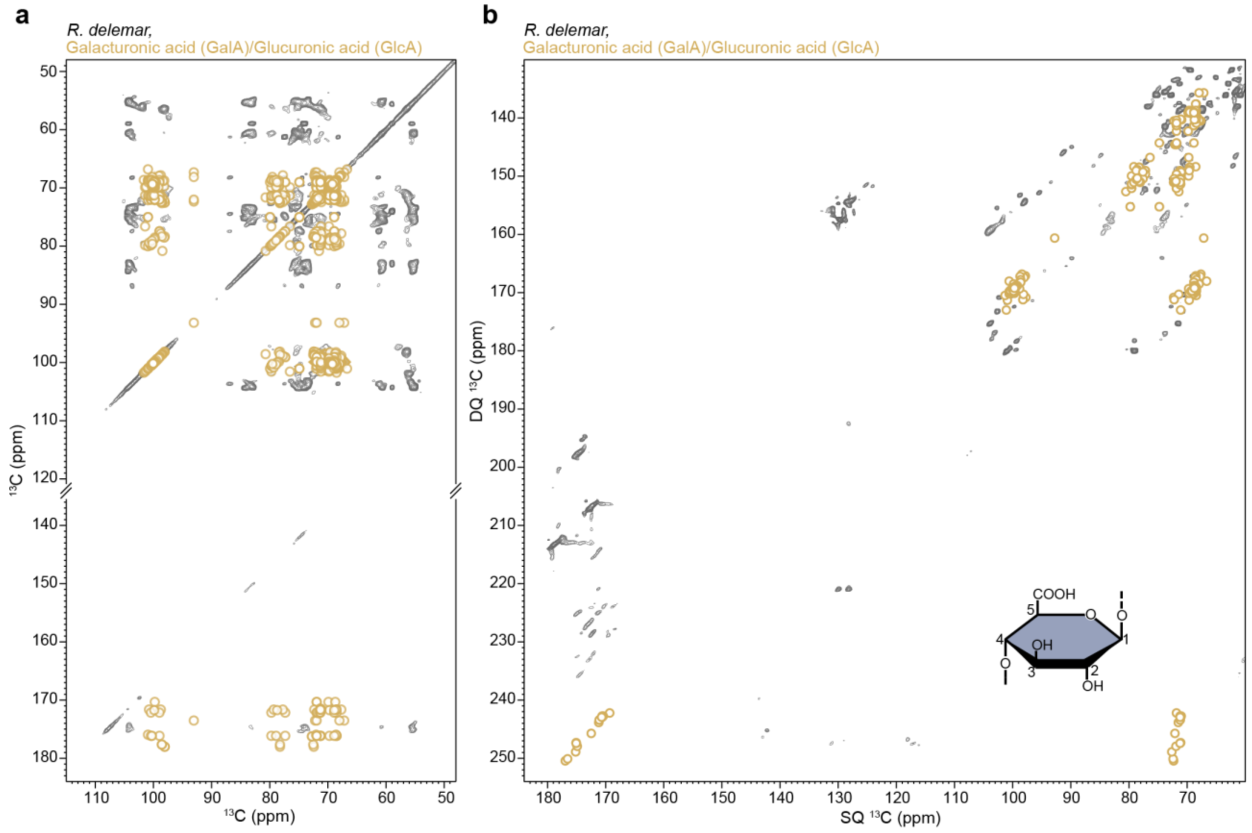


**Supplementary Figure 8. Absence of galacturonic acid and glucuronic acid in *R. delemar* samples**. Overlay of experimentally measured spectra (plotted in grey) with simulated 2D spectra (plotted in yellow) constructed using CCMRD-deposited chemical shifts of galacturonic acid or glucuronic acid. Galacturonic acid and glucuronic acid are not observed within (**a**) 53-ms $^{13}$C-$^{13}$C CORD spectrum or (**b**) $^{13}$C DP J-INADEQUATE; therefore, they are not present in either the rigid or the mobile portions of the cell wall. A representative structure of an GlcA residue is shown.

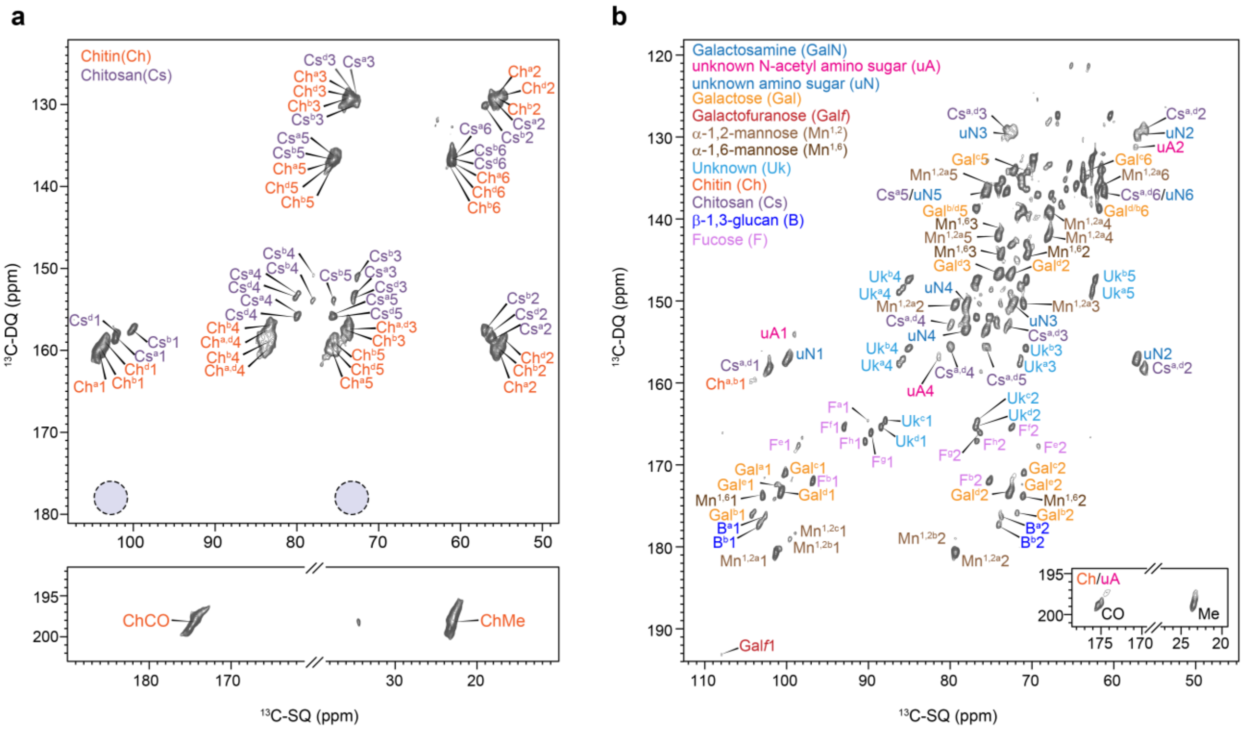


**Supplementary Figure 9. β-glucans in the mobile domain of nikkomycin-treated *R. delemar* cell walls.** (**a**) CP-based 2D $^{13}$C refocused J-INADEQUATE spectrum detecting rigid molecules. β-glucan is absent as highlighted by the dashed line circles. (**b**) Mobile carbohydrates detected by DP-based 2D $^{13}$C refocused J-INADEQUATE spectrum, with signals from β-1,3-glucan (abbreviated as B). Each peak is annotated with the abbreviation of the carbohydrate name, the subtype (in superscript), and the carbon number.

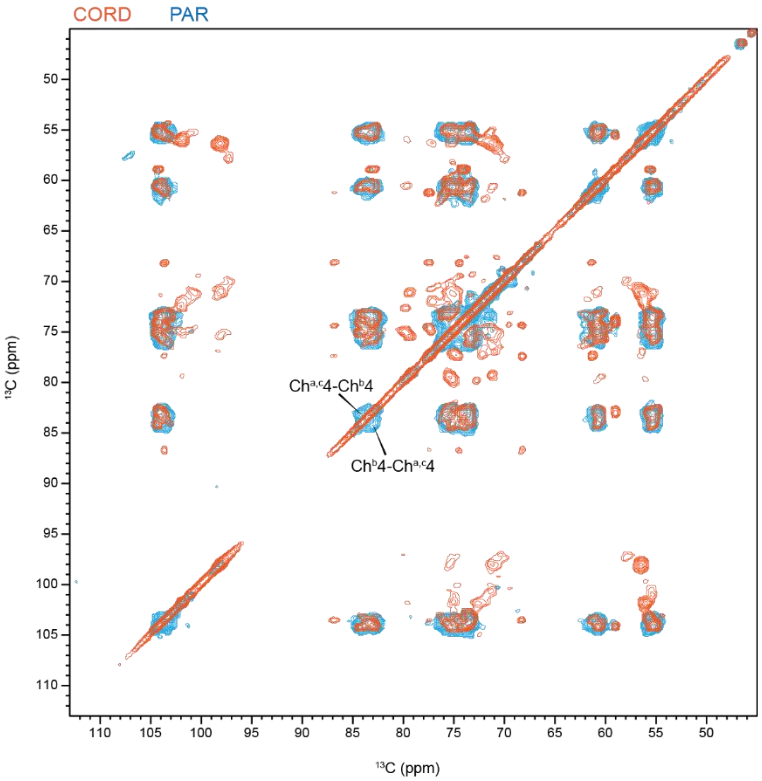


**Supplementary Figure 10. Chitin is the only carbohydrate detectable in 15 ms PAR spectrum.** The overlay of 53 ms CORD spectrum (orange) and 15 ms PAR spectrum (cyan) of *R. delemar*, apo revealed that only chitin signals are retained in both spectra. The signals from β-glucan and chitosan disappeared in PAR, revealing the disordered nature of these two types of molecules. Two cross peaks are observed between different types of chitin molecules in the PAR spectra.

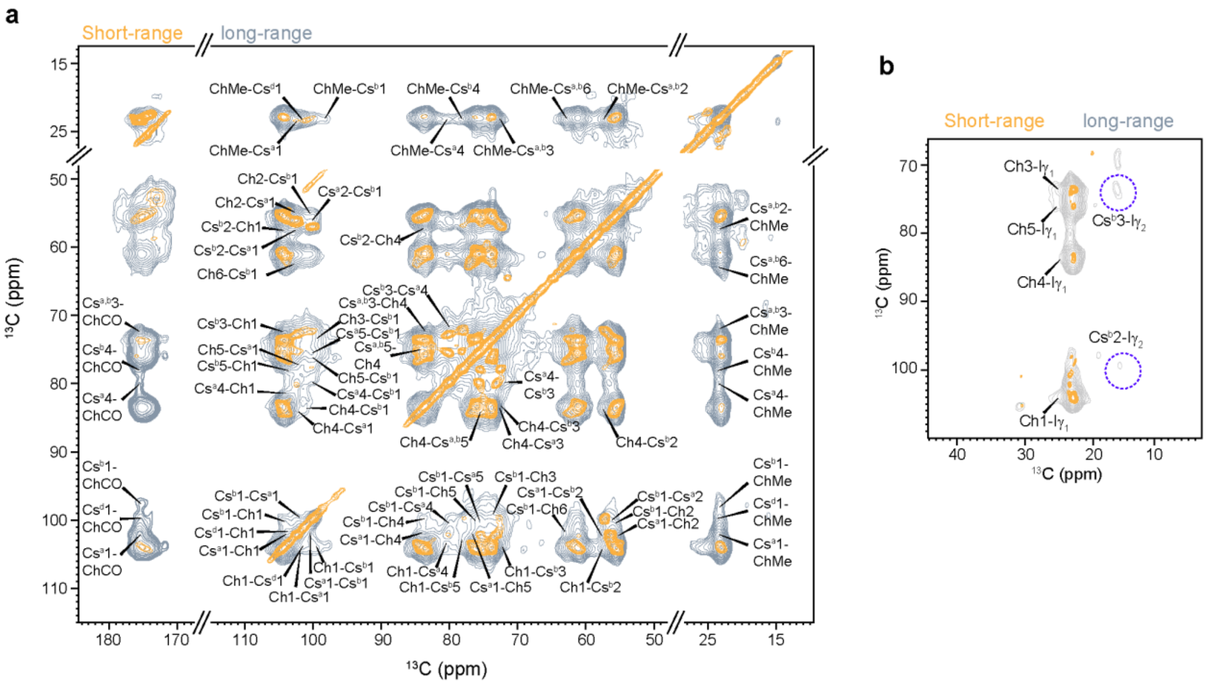

**Supplementary Figure 11. Intermolecular contacts observed in nikkomycin-treated *R. delemar*.** (**a**) Overlay of 2D $^{13}C$-$^{13}C$ 1-s PDSD (grey) with 2D 53-ms CORD spectra (yellow) measured on nikkomycin-treated *R. delemar* cells an 800 MHz spectrometer at 273 K (1s PDSD) or 290 K (53ms CORD) under 15 kHz MAS. Intermolecular cross peaks among rigid polysaccharides are labeled. (**b**) Spectral region (CORD in yellow and PDSD in grey) showing spatial contacts between protein and polysaccharides where the corresponding cross peaks are identified for chitin-isoleucine and chitosan-isoleucine.

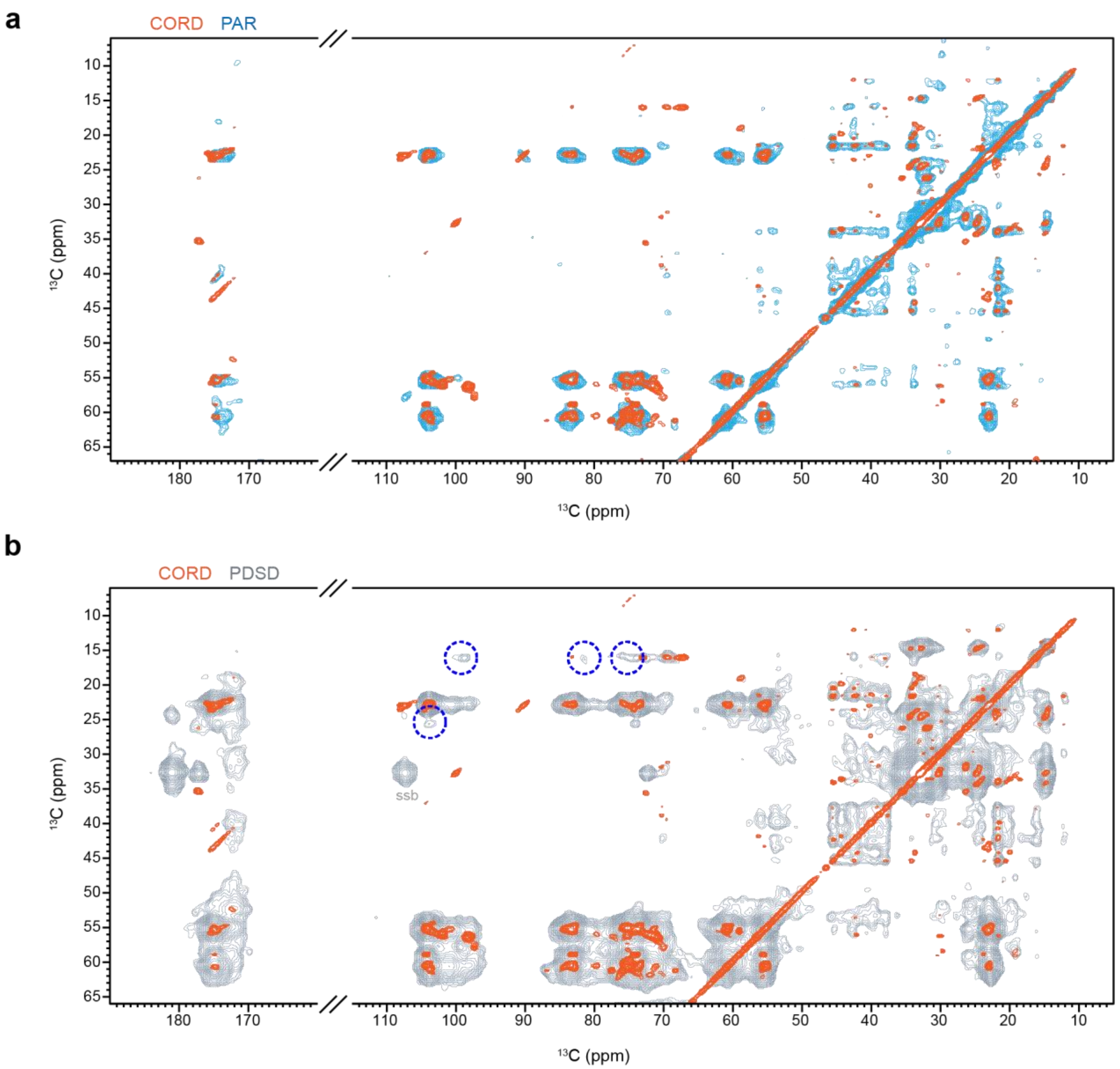


**Supplementary Figure 12. Proteins are partially ordered and closely packed in *R. delemar*, apo. (a)** Overlay of 2D $^{13}C$-$^{13}C$ correlation spectra measured with 53 ms CORD (orange) and 15 ms PAR (cyan). **(b)** Overlay of 2D spectra measured with 53 ms CORD (orange) and 1 s PDSD (grey). Proteins showed largely equilibrated signals in both PAR and PDSD spectra.

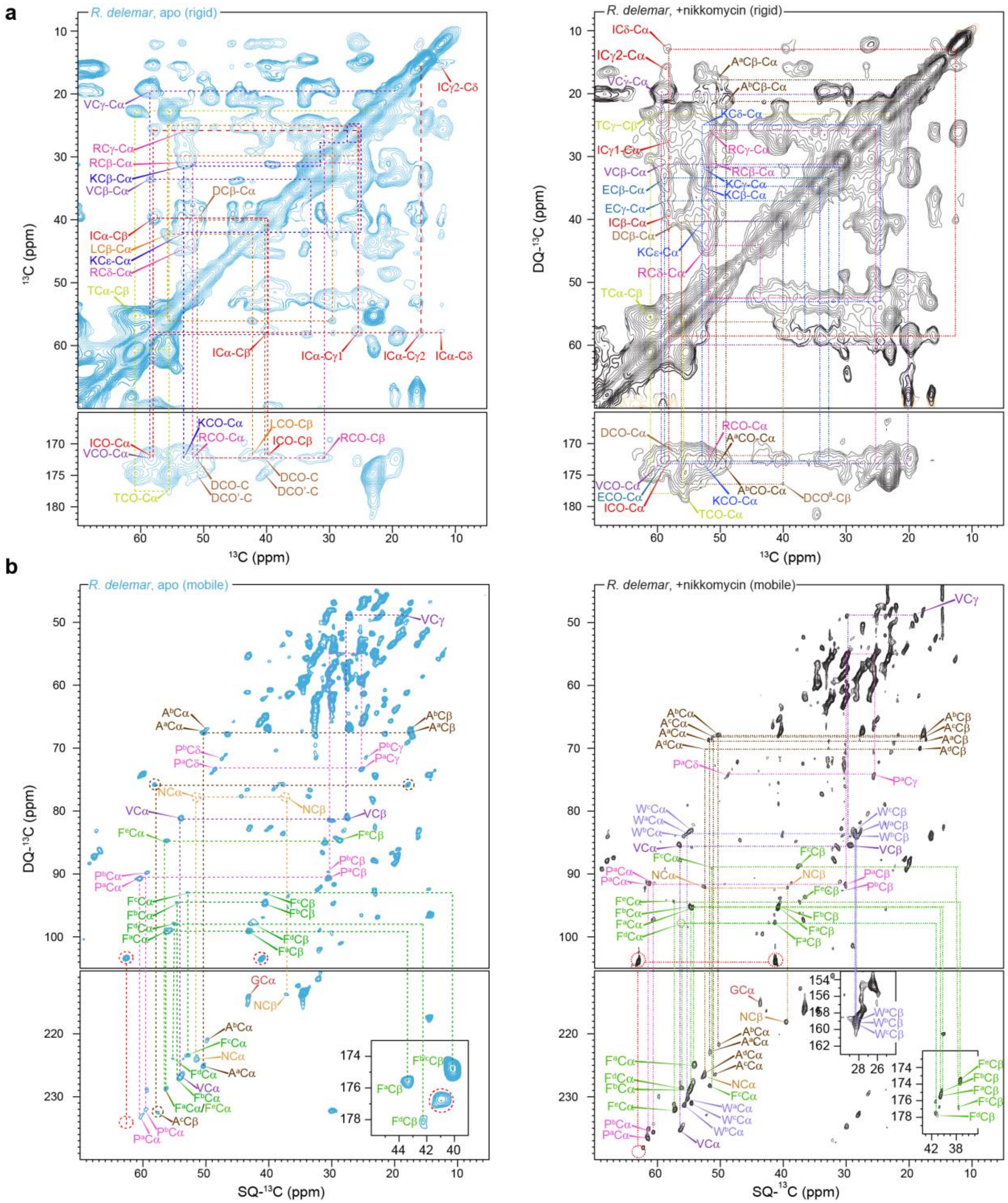

**Supplementary Figure 13. Presence of protein in both mobile and rigid domains.** (**a**) 2D $^{13}C$-$^{13}C$ CP-based 53 ms CORD spectra of *R. delemar* samples (left, apo; right, nikkomycin-treated) showing signals of proteins in the rigid domains. Both spectra were measured on an 800 MHz spectrometer and were processed using Gaussian multiplication (GM) with LB of -30 Hz and GB of 0.03 for both dimensions (F1 and F2). This processing enhanced sensitivity to reveal otherwise weak protein signals. (**b**) 2D $^{13}C$ DP refocused J-INADEQUATE spectra detecting mobile proteins. Single-letter abbreviations are used for the identified amino acids.

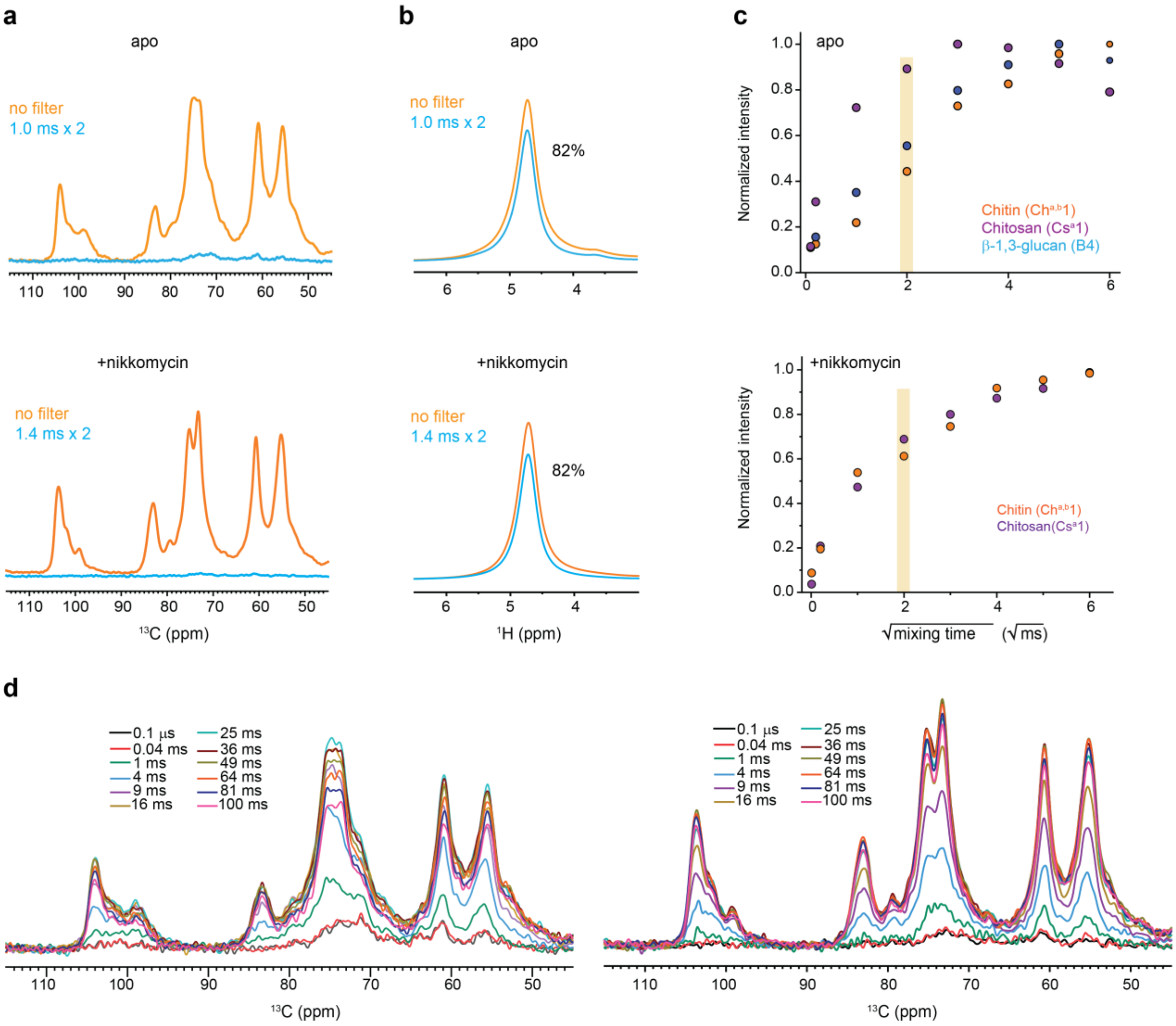


**Supplementary Figure 14. Water-edited experiment setup for inspecting carbohydrate hydration.** (**a**) $^{1}$H-T$_2$ filtered (blue) and control (orange) $^{13}$C spectra are shown for apo (top) and nikkomycin-treated (bottom) *R. delemar* samples. No spin diffusion was applied. Approximately 95% of carbohydrate $^{13}$C signals were removed by the T$_2$ filter. (**b**) $^{1}$H-T$_2$ filtered (blue) and control (orange) $^{1}$H NMR spectra, with 82% of water signal retained for both samples after the $^{1}$H T$_2$ filter. (**c**) Representative water-to-polysaccharide $^{1}$H spin diffusion buildup curves. (**d**) 1D water-edited $^{13}$C spectra with different $^{1}$H mixing times are shown for apo and nikkomycin-treated *R. delemar* samples. All spectra were measured on 400 MHz spectrometer at 15 kHz MAS at 280 K.

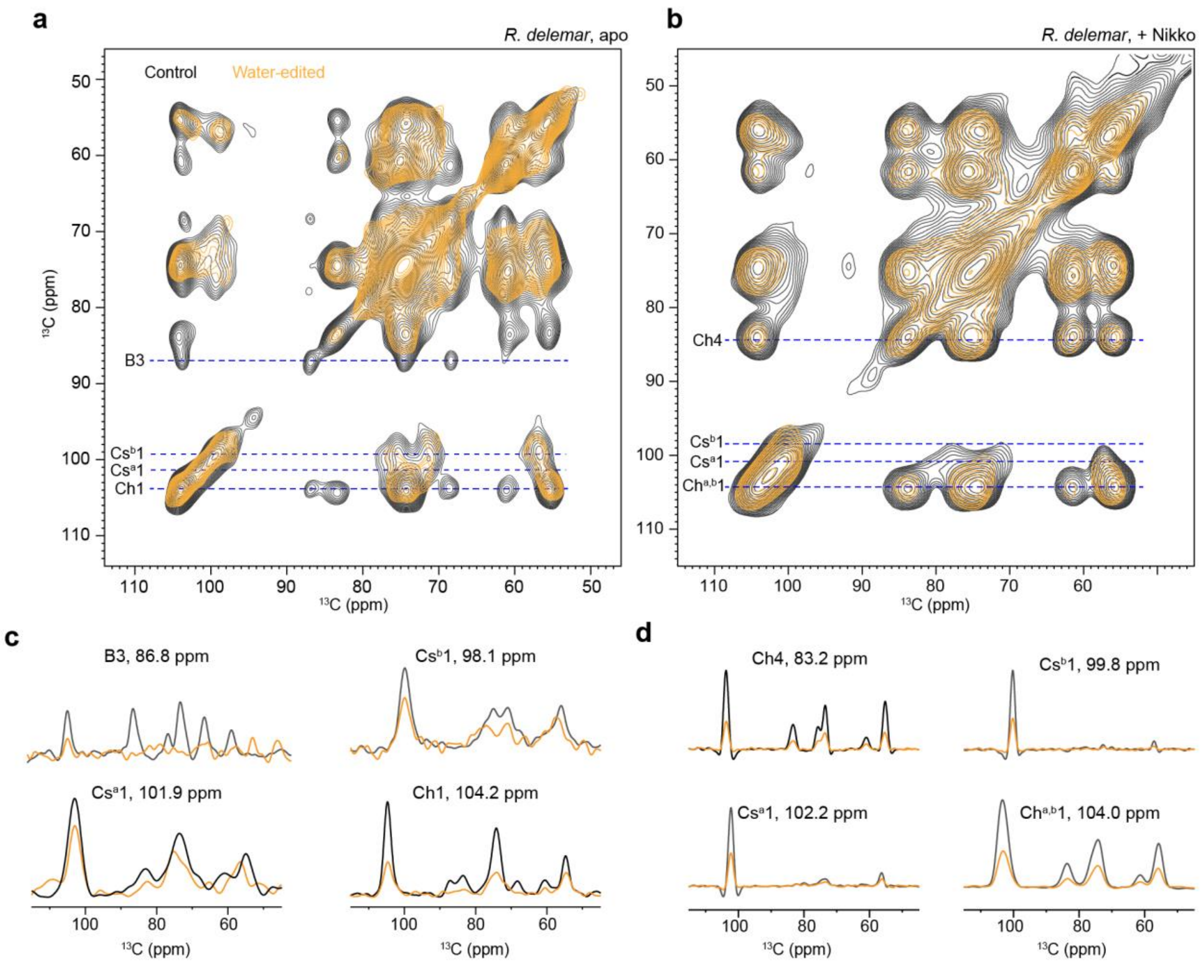


**Supplementary Figure 15. Nikkomycin alters water accessibility of rigid polysaccharides.** (**a**) and (**b**) Overlay of 2D water-edited (orange) and control (black) $^{13}C$-$^{13}C$ correlation spectra ($T_2$ = 1.0 ms x 2 for apo and $T_2$=1.4 ms × 2 for nikkomycin-treated *R. delemar* and $\tau_{SD}$ = 4 ms) for both *R. delemar* and nikkomycin-treated *R. delemar*. The spectra were collected on a 400 MHz NMR spectrometer under 15 kHz MAS at 280 K. Representative 1D slices extracted from the 2D $^{13}C$-$^{13}C$ correlation spectra are shown for (**c**) apo *R. delemar* sample and (**d**) nikkomycin-treated *R. delemar* sample. The control data are displayed as black solid lines, and the water-edited spectra are plotted in orange. All spectra were processed with Gaussian multiplication window function with -30 Hz LB and 0.03 GB (applied in both F1 and F2 dimensions).

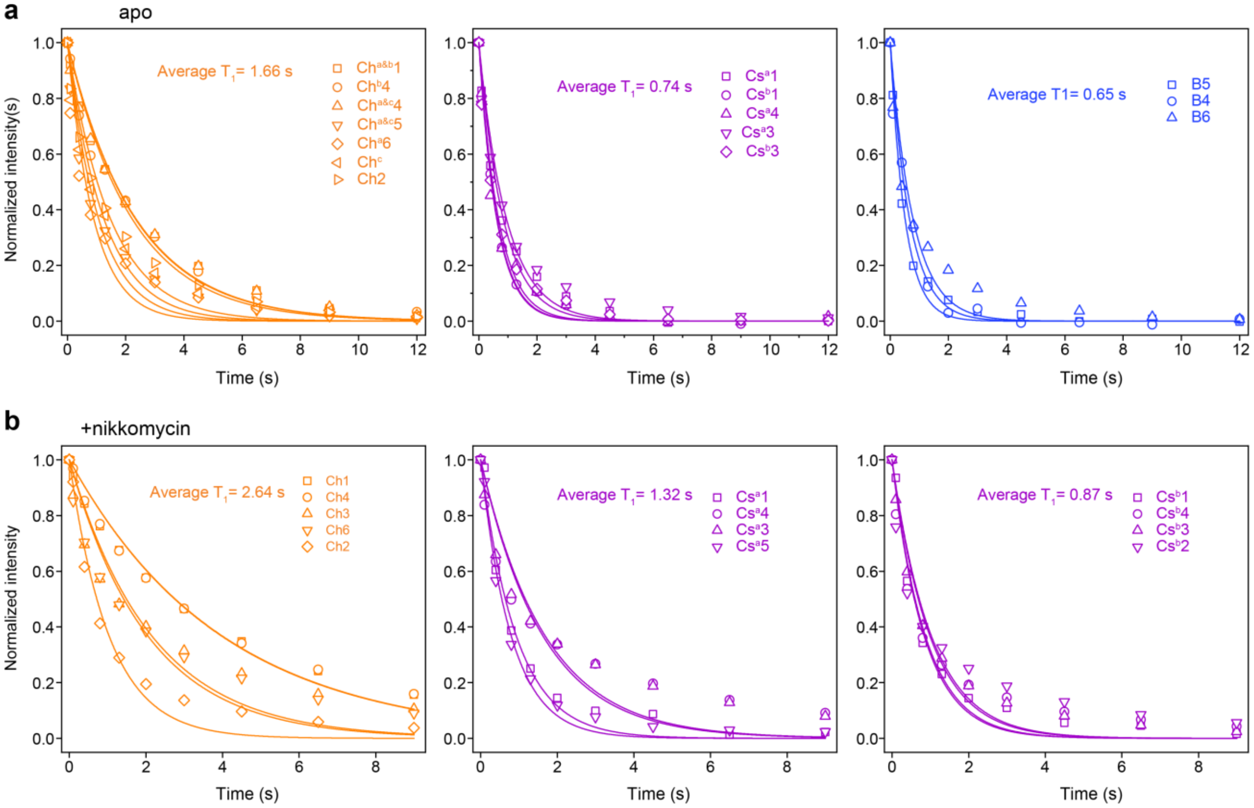


**Supplementary Figure 16. $^{13}C$-$T_1$ of polysaccharides in apo and nikkomycin-treated *R. delemar*.** $^{13}C$-$T_1$ measured with Torchia CP for (**a**) apo and (**b**) nikkomycin-treated samples. The data are separately presented for chitin (orange), chitosan (purple), and β-1,3-glucan (blue). The acquired data were fitted to a single exponential decay equation. Different symbols and color codes are used to represent different carbons in these polysaccharides.

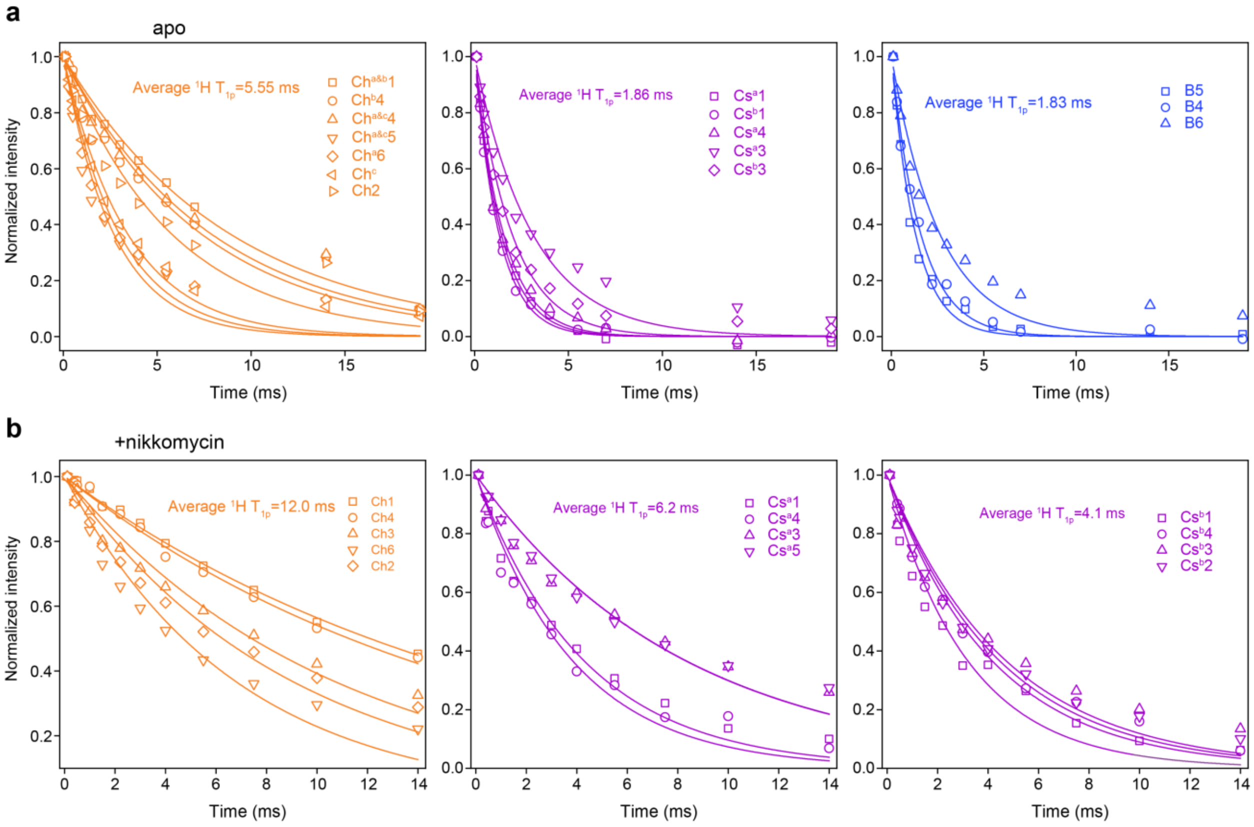


**Supplementary Figure 17. $^{1}$H-T$_{1\rho}$ of polysaccharides in *R. delemar* samples.** $^{1}$H-T$_{1\rho}$ relaxation curves of (**a**) apo and (**b**) nikkomycin-treated *R. delemar* cell walls. The data are separately presented for chitin (orange), chitosan (purple), and β-1,3-glucan (blue). The symbolic representation of each resolvable carbon site is provided.

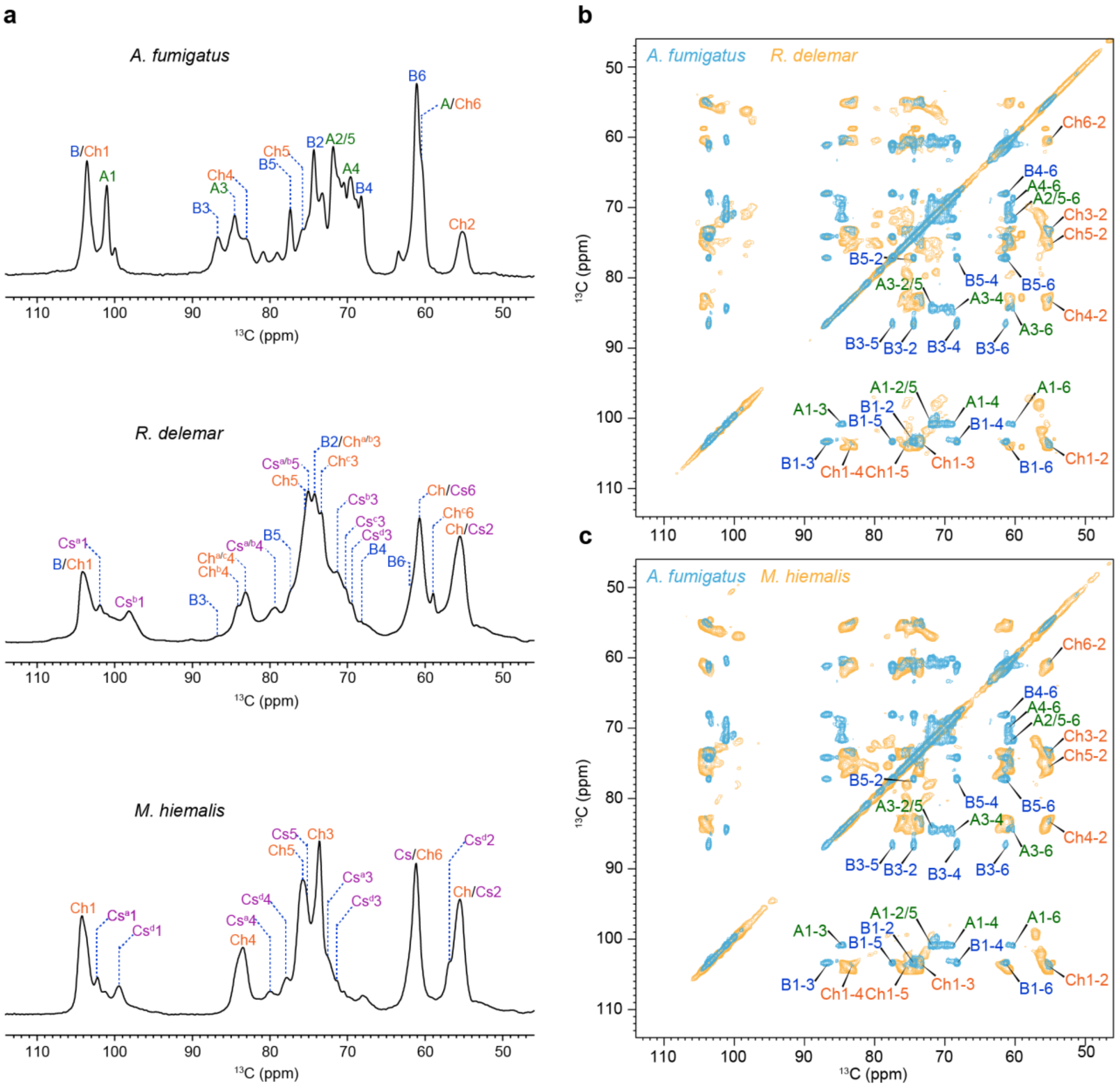


**Supplementary Figure 18. Distinct spectral patterns of *Rhizopus*, *Mucor*, and *Aspergillus* species.** (**a**) Comparison of 1D $^{13}$C CP spectra for *A. fumigatus* (top), *R. delemar* (middle), and *M. hiemalis* (bottom). (**b**) Overlay of 2D $^{13}$C CORD spectra measured on *A. fumigatus* (cyan) and *R. delemar* (orange). (**c**) Overlay of 2D $^{13}$C CORD spectra measured on *A. fumigatus* (cyan) and *M. hiemalis* (orange). The spectra of *A. fumigatus* were adapted from Chakraborty et al.,[1] which is an open access publication. Resonance assignments of *A. fumigatus* polysaccharides were shown for 2D spectra. All spectra were collected on 800 MHz NMR spectrometers.

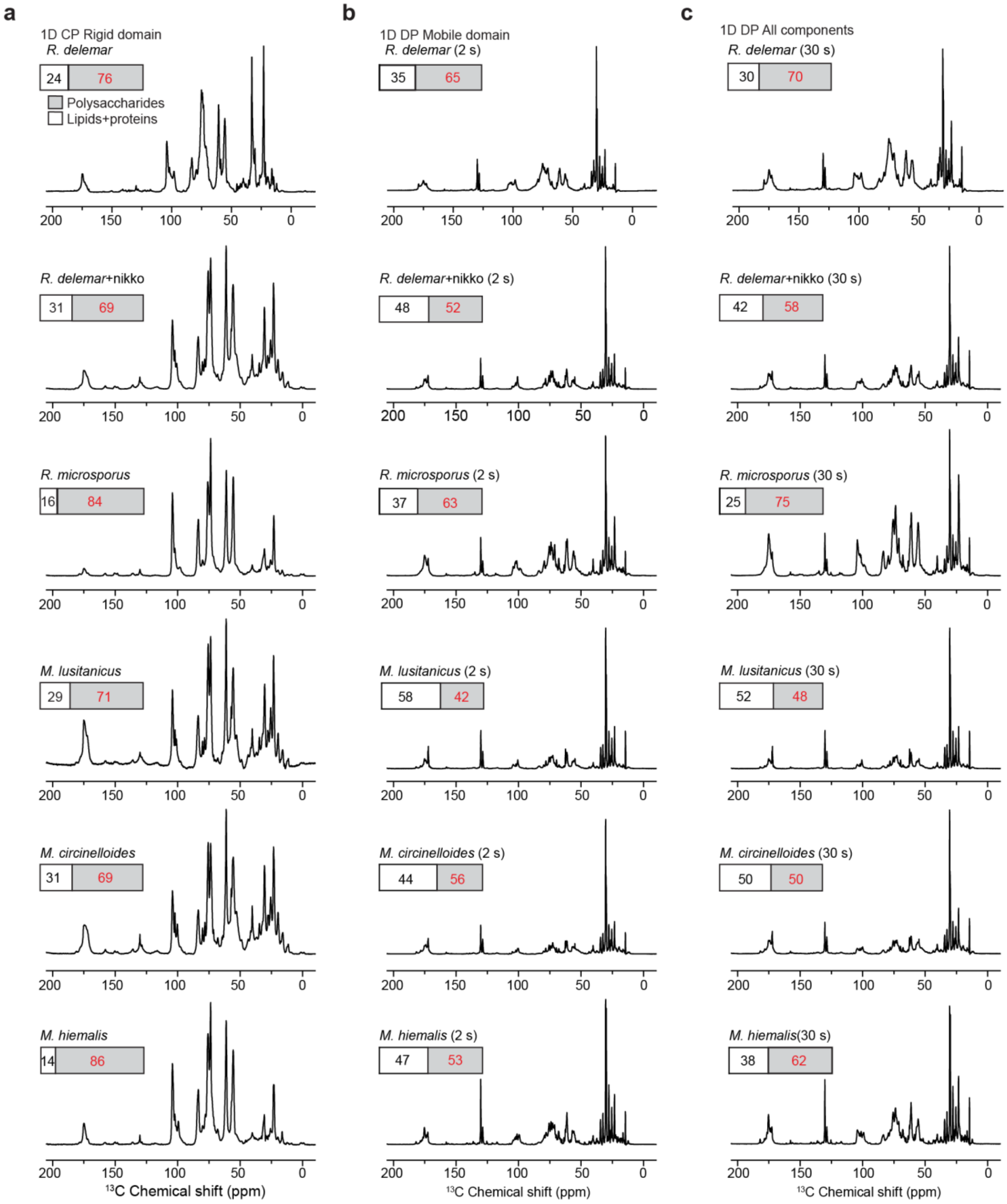


**Supplementary Figure 19. Similarity of *Rhizopus* and *Mucor* carbohydrates.** (**a**) 1D $^{13}$C CP spectra for the detection of the rigid components within fungal cells. (**b**) and (**c**) 1D $^{13}$C DP spectra measured with both short recycle delays of 2 s (mainly detects mobile components) and long recycle delays of 30 s (enabling quantitative detection for all components of the intact fungal cells).

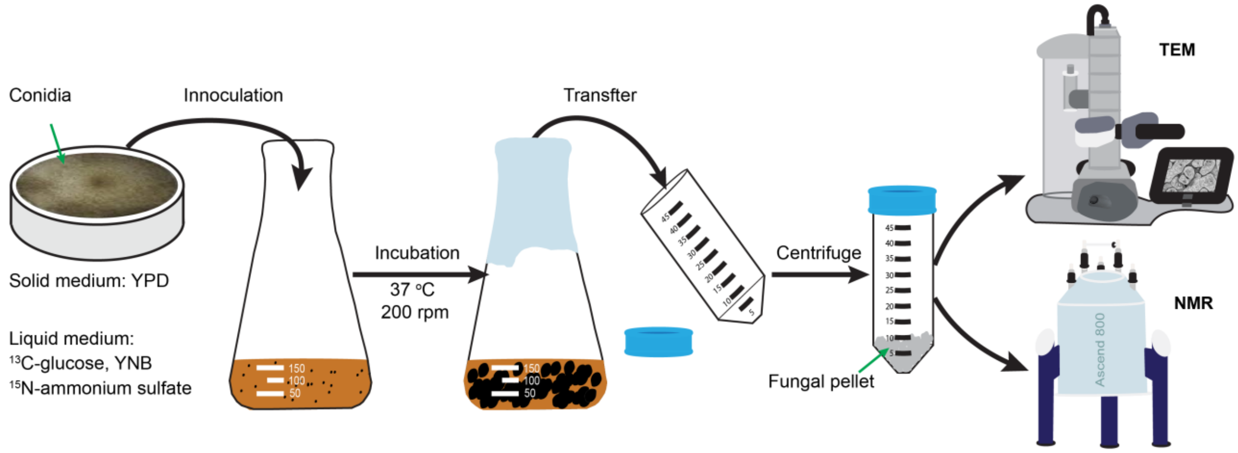


**Supplementary Figure 20. Experimental flowchart of fungal cultivation.** Initially, fungal conidia cultivated on YPD solid medium are inoculated into a 250-mL shake flask pre-filled with 100 ml of sterilized growth medium ($^{13}$C-glucose, $^{15}$N-ammonium sulfate, and YNB). Subsequently, the shake flask is placed in a shaker for 5 days, maintaining a temperature of 37 °C and a shaking rate of 200 rpm. Following cultivation, the solution in the shake flask is transferred to 50-mL centrifuge flasks and centrifuged for 20 minutes at 6,500 rpm at 4 °C to collect the fungal pellets. The fungal pellets are thoroughly washed with phosphate-buffered saline (PBS) to eliminate any trace amounts of $^{13}$C- or $^{15}$N-labeled water-soluble fungal metabolites or source materials from the growth medium. The collected fungal pellets are stored in a freezer at -20 °C before TEM imaging or NMR measurements.

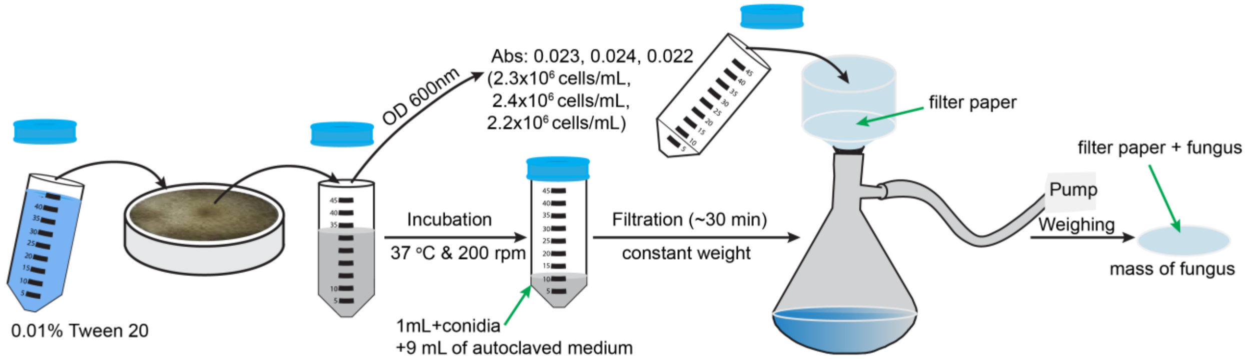


**Supplementary Figure 21. Experimental flowchart of constructing fungal growth curves.** First, fungal conidia are harvested using 0.01% Tween-20, and the optical density (OD) at 600 nm (with 0.01% Tween-20 solution) is measured. Second, 1 mL of the fungal conidia solution is transferred into a sterile 50-mL conical tube containing 9 mL of autoclaved growth medium. The tube is then placed in a cultivation shaker at 37 °C and 200 rpm. At designated time intervals of the cultivation period, the 50-mL conical tube is removed, and pump filtration is conducted for approximately 30 min to achieve a constant weight. Mass is determined at each specific time point during cultivation. All measurements are triplicated.

**Supplementary Table S1. Measurements of the cell-wall thickness of *R. delemar*.** All measurements are measured by image J and the unit is nm.[2] Reading x is abbreviated as Rx, e.g., R1 for Reading 1.

| *R. delemar*, apo | | | | | | | | | | |
|---|---|---|---|---|---|---|---|---|---|---|
| Reading | Cell 1 | Cell 2 | Cell 3 | Cell 4 | Cell 5 | Cell 6 | Cell 7 | Cell 8 | Cell 9 | Cell 10 |
| R1 | 129 | 118 | 156 | 146 | 116 | 101 | 113 | 132 | 101 | 143 |
| R2 | 116 | 121 | 157 | 137 | 110 | 102 | 110 | 152 | 124 | 108 |
| R3 | 100 | 117 | 125 | 139 | 123 | 105 | 117 | 131 | 102 | 104 |
| R4 | 107 | 135 | 119 | 141 | 123 | 110 | 122 | 134 | 119 | 137 |
| R5 | 110 | 131 | 139 | 121 | 104 | 97 | 135 | 95 | 94 | 155 |
| R6 | 98 | 138 | 147 | 135 | 124 | 104 | 120 | 101 | 96 | 124 |
| R7 | 102 | 131 | 129 | 119 | 138 | 122 | 109 | 108 | 106 | 132 |
| R8 | 120 | 112 | 103 | 115 | 136 | 101 | 113 | 89 | 112 | 136 |
| R9 | 104 | 102 | 116 | 144 | 111 | 100 | 135 | 104 | 109 | 134 |
| R10 | 140 | 118 | 103 | 121 | 126 | 94 | 129 | 94 | 102 | 122 |
| Average | 113±13 | 122±11 | 129±19 | 132±11 | 121±10 | 104±7 | 122±9 | 114±20 | 107±9 | 129±14 |
| Average for all readings | 119±16 | | | | | | | | | |
| *R. delemar*, + nikkomycin | | | | | | | | | | |
| R1 | 226 | 181 | 173 | 197 | 196 | 191 | 251 | 246 | 265 | 224 |
| R2 | 234 | 192 | 186 | 197 | 183 | 197 | 259 | 241 | 274 | 224 |
| R3 | 226 | 176 | 168 | 213 | 202 | 183 | 214 | 237 | 268 | 212 |
| R4 | 207 | 171 | 173 | 224 | 196 | 191 | 218 | 237 | 274 | 239 |
| R5 | 194 | 172 | 192 | 208 | 205 | 195 | 236 | 250 | 276 | 228 |
| R6 | 216 | 190 | 179 | 194 | 172 | 202 | 222 | 220 | 271 | 243 |
| R7 | 231 | 171 | 185 | 188 | 183 | 188 | 235 | 257 | 262 | 228 |
| R8 | 233 | 215 | 168 | 189 | 164 | 169 | 229 | 239 | 253 | 225 |
| R9 | 216 | 202 | 196 | 176 | 199 | 192 | 251 | 239 | 241 | 222 |
| R10 | 200 | 176 | 180 | 203 | 198 | 190 | 252 | 251 | 278 | 214 |
| Average | 218±13 | 184±14 | 179±9 | 198±13 | 189±13 | 189±9 | 237±15 | 241±10 | 266±11 | 226±9 |
| Average for all readings | 213±27 | | | | | | | | | |

**Supplementary Table S2. Solid-state NMR experimental parameters for fungal cell wall characterization.** T = sample temperature; $B_0$ = magnetic field; $\nu_{MAS}$ = MAS frequency; ns = number of scans; $d_1$ = recycle delay between scans; $t_{1, max}$ = maximum $t_1$ evolution time (for indirect dimension); $t_{1, inc}$ = increment for $t_1$ (for indirect dimension) evolution time; $\tau_{dw}$ = dwell time during direct FID acquisition; $\tau_{acq}$ = maximum acquisition time during direct FID detection; $\tau_{XY}$ = cross-polarization contact time during CP from channel X to channel Y; $\nu_{1H, dec}$ = dipolar decoupling field strength.

| Experiment | T (K) | $B_0$ (T) | $\nu_{MAS}$ (kHz) | NS | d1 (s) | $t_{1, max}$ (ms) | $t_{1, inc}$ (μs) | $\tau_{dw}$ (μs) | $\tau_{acq}$ (ms) | $\tau_{HC}$ (ms) | $\tau_{mix}$ (ms) | $\nu_{1H\ dec}$ (kHz) | Samples |
|---|---|---|---|---|---|---|---|---|---|---|---|---|---|
| 1D $^{13}C$ CP | 290 | 18.8 | 13.5 or 15 [a] | 256 or 1024 | 2 | | | 7/5 | 19.6/18 | 1 | | 83 or 83 | *R. delemar*, apo/+nikko or other mucor fungi |
| 1D $^{13}C$ DP | 290 | 18.8 | 13.5 or 15 | 1024 | 2 | | | 7/5 | 28.7/18 | | | 100 or 83 | |
| 1D $^{13}C$ DP | 290 | 18.8 | 13.5 or 15 | 96 or 192 | 30 or 35 | | | | | | | | |
| 2D CORD | 290 | 18.8 | 13.5 or 15 | 16 | 1.6 | 7 | 25 | 7.5 or 5 | 18/14 | 1 | 53 $\tau_{CORD}$ | 83 or 71 | |
| 2D CP J-INADEQUATE | 290 | 18.8 | 13.5 or 15 | 16 | 1.5 or 2 | 8 or 7.5 | 60 or 22 | 7.5 or 5 | 18/14 | 0.5 | | 92.5 or 71 | *R. delemar*, apo and nikkomycin-treated |
| 2D DP J-INADEQUATE | 290 | 18.8 | 13.5 or 15 | 16 | 1.5 or 2 | 10 or 7.5 | 20 or 22 | 7.5 or 5 | 19/14 | | | 83 | |
| 2D water-edited | 280 | 9.4 | 15 | 256 or 64 | 2 | 5.5 | 50 | 8 | 16 | 1 | 50 $\tau_{PDSD}$ | 83 | |
| 1D Torchia $^{13}C$-$T_1$ | 298 | 9.4 | 10 or 15 | 256 | 2 | | | 10/8 | 14/16 | 1 | 50 $\tau_{PDSD}$ | 71 or 83 | |
| 2D PDSD | 273 | 18.8 | 15 | 64 | 2 | 6.5 | 25 | 5 | 14 | 1 | 1000 $\tau_{PDSD}$ | 83 | |
| 2D PAR | 290 | 18.8 | 13.5 | 64 | 1.7 | 5.7 | 27 | 7.5 | 16 | 0.5 | 15 $\tau_{PAR}$ | 83 | *R. delemar*, apo |
| 2D $^{15}N$-$^{1}H$ HETCOR | 290 | 18.8 | 13.5 | 8 | 1.6 | 2.3 | 41 | 7.5 | 18 | | | 92.5 | |
| 2D N(CA)CX with DARR | 290 | 18.8 | 13.5 | 32 | 1.7 | 10 | 100 | 7.5 | 18 | | 100 $\tau_{DARR}$ | 92.5 | |
| 2D hCH | 302 | 18.8 | 60 | 16 | 2 | 4.26 | 33.33 | 8.5 | 19.9 | 0.2 for CP1&2 | | 10 | |
| 2D hNH | 302 | 18.8 | 60 | 16 | 2 | 6.4 | 50.12 | 15.3 | 19.9 | 2 for CP1, 0.2 or 2 for CP2 | | 10 | |

[a] 13.5 kHz for *R. delemar* apo sample except 1s-PDSD (15 kHz and 273 K) and 2D water-edited experiments (15 kHz and 280 K). 15 kHz for nikkomycin-treated *R. delemar* sample and other *Rhizopus* and *Mucor* species.

**Supplementary Table S3. $^{13}C$ and $^{15}N$ chemical shifts of rigid polysaccharides in *R. delemar* cell walls**. Superscripts are used to denote different allomorphs. Not applicable (/). Unidentified (-). Minor forms (m).

| Carbohydrate | | C1 | C2 | C3 | C4 | C5 | C6 | CO | $CH_3$ | N | Experiments | References |
|---|---|---|---|---|---|---|---|---|---|---|---|---|
| ***R. delemar*, apo** | | | | | | | | | | | | |
| chitin | a | 104.2 | 55.3 | 74.0 | 83.3 | 75.7 | 60.5 | 174.5 | 22.9/22.7 | 123.8 | $^{13}C$-$^{13}C$ CORD, $^{13}C$ CP J-INADEQUATE | Kang et al., 2018 and Fernando et al., 2021[3-4] |
| | b | 104.3 | 55.3 | 74.1 | 84.3 | 75.1 | 60.7 | 174.9 | 22.9 | 123.7 | | |
| | c | 104.3 | 55.4 | 74.0 | 83.1 | 75.1 | 58.9 | 174.9 | 23.7 | 123.9 | | |
| | d(m) | 104.3 | 54.9 | 73.2 | 83.4 | 76.2 | 60.9 | 173.1 | 22.4 | 123.6 | | |
| chitosan | a | 101.9 | 56.0 | 72.6 | 79.5 | 75.0 | 60.7/60.3 | / | / | 32.6 | | |
| | b | 98.1 | 56.5 | 71.1 | 79.8/77.0 | 75.3 | 60.7 | / | / | 33.6 | | |
| | c(m) | 97.3 | 57.8 | 70.5 | 80.0 | 75.2 | 60.8 | / | / | 33.6 | | |
| | d | 100.7 | 55.4 | 70.0 | 78.8 | 74.7 | 61.6 | / | / | 33.6 | | |
| β-1,3-glucan | | 103.7 | 74.4 | 86.8 | 68.2 | 77.4 | 61.3 | / | / | / | | |
| ***R. delemar*, + nikkomycin** | | | | | | | | | | | | |
| chitin | a | 104.4 | 55.2 | 73.6 | 83.2 | 76.3 | 60.5 | 173.9 | 22.8 | - | $^{13}C$-$^{13}C$ CORD, $^{13}C$ CP J-INADEQUATE | Chakraborty et al., 2021[1] |
| | b | 104.0 | 55.6 | 73.9 | 83.5 | 76.4 | 61.1 | 174.9 | 22.9 | - | | |
| | d | 103.7 | 54.9 | 73.5 | 83.4 | 76.0 | 60.8 | 174.6 | 23.1 | - | | |
| chitosan | a | 102.2 | 56.1 | 72.8 | 79.9 | 75.3 | 60.8 | / | / | - | | |
| | d(m) | 101.6 | 56.0 | 72.7 | 80.5 | 75.4 | 61.6 | / | / | - | | |
| | b | 99.8 | 57.0 | 72.4 | 77.6 | 75.6 | 61.0 | / | / | - | | |

**Supplementary Table S4. $^{13}$C chemical shifts of mobile polysaccharides in *R. delemar* cell walls**. Superscripts are used to denote different allomorphs. Not applicable (/). Unidentified (-).

| ***R. delemar*, apo** | | | | | | | |
|---|---|---|---|---|---|---|---|
| Carbohydrate | | C1 | C2 | C3 | C4 | C5 | C6 |
| Ch | a&b | 104.1 | 55.3 | 73.4 | 83.1 | 74.7 | 60.1 |
| | d | 104.3 | 54.7 | 73.2 | 82.9 | 74.9 | 61.0 |
| Cs | a | 101.8 | 56.5 | 72.6 | 77.9 | 75.3 | 60.8 |
| | b | 98.0 | 56.6 | 72.6 | 77.9 | 75.3 | 60.8 |
| GalNAc | | 95.4 | 57.4 | 72.4 | 82.4 | 75.9 | 61.1 |
| GalN | | 91.3 | 54.5 | 69.2 | 77.9 | - | - |
| Mn$^{1,2}$ | a | 101.0 | 78.9 | 70.7 | 67.5 | 73.8 | 61.7 |
| | b | 99.3 | 80.4 | - | - | - | - |
| | c | 99.3 | 79.0 | - | - | - | - |
| Mn$^{1,6}$ | | 102.3 | 70.6 | 73.7 | 67.6 | 71.2 | 66.2 |
| β-1,3-glucan | a | 103.3 | 73.9 | - | - | - | - |
| | b | 103.6 | 74.5 | - | - | - | - |
| α-Fucose | a | 90.0 | 74.3 | 71.1 | 71.1 | 67.7 | 19.6 |
| | b | 96.7 | 74.8 | 77.1 | 70.2 | 68.4 | 20.2 |
| | c | 91.0 | 74.5 | 70.2 | 73.3 | 66.6 | 20.1 |
| | d | 94.5 | 71.9 | 72.3 | 69.7 | 67.1 | 19.2 |
| | e | 98.3 | 67.0 | 70.3 | 71.4 | 67.4 | 15.9 |
| Gal | a | 102.5 | / | / | / | / | / |
| | b | 103.9 | 71.0 | 72.8 | 70.3 | 76.7 | 61.3 |
| | c | 99.9 | 70.7 | 73.2 | 70.1 | 71.5 | 63.3 |
| | d | 99.7 | 72.6 | / | / | / | / |
| | e | 101.0 | 70.6 | / | / | / | / |
| Gal*f* | | 107.3 | 90.5 | / | / | / | / |
| Unknown | a | - | - | 71.3 | 85.9 | 62.4 | - |
| | b | - | - | 70.0 | 85.0 | 61.5 | - |
| uA* | a | 102.7 | 52.7 | - | - | - | - |
| | b | 101.5 | 55.6 | 74.4 | 83.9 | - | - |
| | c | 89.8 | 55.2 | 73.4 | 83.2 | 75.4 | 60.7 |
| uN* | a | 98.8 | 54.3 | 69.2 | 77.9 | - | - |
| | b | 93.4 | 57.5 | 74.6 | 80.0 | - | - |

| ***R. delemar*, + Nikkomycin** | | | | | | | | |
|---|---|---|---|---|---|---|---|---|
| Carbohydrate | | C1 | C2 | C3 | C4 | C5 | C6 | Reference |
| Ch | a&b | 104.2 | 55.5 | - | - | - | - | Chakraborty et al., 2021[1] |
| Cs | a | 102.2 | 56.0 | 72.8 | 79.8 | 75.7 | 61.3 | |
| | d | 99.8 | 57.1 | 72.5 | 78.0 | 75.6 | 61.1 | |
| Mn$^{1,2}$ | a | 101.3 | 79.3 | 71.0 | 67.8 | 74.1 | 62.0 | |
| | b | 99.6 | 79.4 | - | - | - | - | |
| | c | 98.9 | 79.4 | - | - | - | - | |
| Mn$^{1,6}$ | | 102.8 | 71.0 | 73.7 | 67.5 | 71.7 | 66.3 | |
| β-1,3-glucan | a | 102.5 | 73.6 | 82.9 | - | - | - | |
| | b | 103.7 | 74.0 | 85.7 | - | - | - | |
| α-Fucose | a | 90.1 | 74.5 | - | - | - | - | Furevi et al., 2022[5] |
| | b | 96.7 | 75.2 | 76.6 | 70.6 | 68.8 | 20.4 | |
| | e | 98.5 | 69.2 | 73.8 | 70.1 | 69.7 | 15.9 | |
| | f | 93.0 | 72.2 | 70.4 | 69.2 | 68.0 | 19.9 | |
| | g | 90.0 | 76.4 | 72.9 | 73.3 | 68.0 | 19.9 | |
| | h | 90.7 | 76.5 | 73.8 | 71.8 | 68.0 | 14.4 | |
| Gal | b | 104.2 | 73.4 | - | - | 76.8 | 61.9 | Kang et al., 2018[4] |
| | c | 100.2 | 70.8 | 68.0 | 74.1 | 70.4 | 63.7 | Fernando et al., 2023[6] |
| | d | 100.6 | 72.7 | 73.7 | 70.6 | 76.9 | 61.7 | |
| | e | 101.4 | 70.1 | - | - | - | - | |
| Galf | | 107.7 | 86.0 | - | - | - | - | |
| Unknown | a | - | - | 70.6 | 85.0 | 62.1 | - | |
| | b | - | - | 71.4 | 86.2 | 62.5 | - | |
| | c | 88.0 | 76.6 | - | - | - | - | |
| | d | 88.5 | 77.1 | - | - | - | - | |
| uN* | | 99.9 | 57.1 | 72.6 | 78.0 | 75.6 | 61.1 | |
| uA* | | 99.1 | 55.0 | - | 81.4 | - | - | |

* Note: Unknown amino sugars without (uN) and with (uA) acetyl groups are categorized empirically by their C4 chemical shifts (> 81 ppm for uA; < 81 ppm for uN).

**Supplementary Table S5. Chemical shifts of amino acids in *R. delemar* cells.** Superscripts are used to represent the same type of amino acids (AAs). Not applicable (/). Unidentified (-). The rigid AAs are identified by 2D 53 ms $^{13}C$-$^{13}C$ CORD (CORD) spectra and the mobile AAs are assigned using the 2D $^{13}C$ DP J-based INADEQAUTE (DP-INADEQUATE).

| ***R. delemar*, apo** | | | | | | | | |
|---|---|---|---|---|---|---|---|---|
| Amino acids | | Cα | Cβ | Cγ | Cδ | Cε | CO | CO’ |
| **Rigid AAs (CORD)** | | | | | | | | |
| Aspartic | D | 50.8 | 40.1 | / | / | / | 171.7 | 174.6 |
| Arginine | R | 52.0 | 30.5 | 25.3 | 44.0 | / | 172.1 | / |
| Isoleucine | I | 58.0 | 40.0 | 25.6(15.5) | 12.4 | / | 171.6 | / |
| Leucine | L | 55.7 | 42.6 | 29.7 | 25.2 | / | 171.7 | / |
| Lysine | K | 53.3 | 30.9 | 28.1 | 25.4 | 42.1 | 172.3 | / |
| Threonine | T | 56.0 | 60.7 | 22.7 | / | / | 174.7 | / |
| Valine | V | 58.1 | 33.1 | 19.1 | / | / | 172.4 | / |
| **Mobile AAs (DP-INADEQUATE)** | | | | | | | | |
| Alanine | $A^{a}$ | 50.2 | 17.4 | / | / | / | 174.9 | / |
| | $A^{b}$ | 49.7 | 17.3 | / | / | / | 174.9 | / |
| Asparagine | N | 51.2 | 37.0 | / | / | / | 172.3 | 176.7 |
| Glycine | G | 43.1 | / | / | / | / | 172.0 | / |
| Phenylalanine | $F^{e}$ | 56.0 | 28.8 | 124.5/128.3 | - | - | 171.2 | / |
| Phenylalanine | $F^{a}$ | 55.9 | 43.0 | 132.1 | - | - | 172.5 | / |
| | $F^{b}$ | 54.3 | 40.3 | 134.6 | - | - | 171.9 | / |
| | $F^{c}$ | 52.9 | 40.2 | 136.1 | - | - | 171.3 | / |
| | $F^{d}$ | 55.0 | 42.8 | 135.9 | - | - | 169.6 | / |
| Proline | $P^{a}$ | 60.4 | 30.3 | 25.2 | 48.5 | / | 172.6 | / |
| | $P^{b}$ | 59.3 | 30.7 | 24.3 | 47.3 | / | 169.2 | / |

| ***R. delemar*, + nikkomycin** | | | | | | | | |
|---|---|---|---|---|---|---|---|---|
| Amino acids | | Cα | Cβ | Cγ | Cδ | Cε | CO | CO’ |
| **Rigid AAs (CORD)** | | | | | | | | |
| Alanine | $A^{a}$ | 51.0 | 17.5 | / | / | / | 173.2 | / |
| | $A^{b}$ | 49.2 | 21.4 | / | / | / | 173.3 | / |
| Arginine | R | 52.3 | 32.2 | 25.7 | 44.1 | / | 172.5 | / |
| Isoleucine | I | 58.3 | 39.5 | 27.0(15.9) | 12.8 | / | 172.9 | / |
| Aspartic | D | 56.1 | 40.3 | / | / | / | 172.6 | 176.9 |
| Lysine | K | 52.9 | 34.9 | 31.5 | 25.0 | 40.2 | 173.0 | / |
| Glutamic | E | 55.9 | 32.9 | 37.2 | / | / | 172.8 | 176.2 |
| Valine | V | 60.2 | 31.1 | 19.8 | / | / | 173.8 | / |
| Threonine | T | 56.1 | 61.4 | 23.3 | / | / | 173.0 | / |
| **Mobile AAs (DP-INADEQUATE)** | | | | | | | | |
| Alanine | $A^{a}$ | 52.0 | 17.3 | / | / | / | 171.4 | / |
| | $A^{b}$ | 50.2 | 17.6 | / | / | / | 171.4 | / |
| | $A^{c}$ | 50.9 | 17.0 | / | / | / | 175.5 | / |
| | $A^{d}$ | 52.4 | 17.8 | / | / | / | 173.6 | / |
| Phenylalanine | $F^{a}$ | 54.4 | 41.0 | 136.2 | - | - | 170.3 | / |
| | $F^{b}$ | 54.8 | 40.6 | 134.8 | - | - | 174.1 | / |
| | $F^{c}$ | 51.3 | 37.6 | 139.6 | - | - | 177.3 | / |
| | $F^{d}$ | 56.6 | 413 | 134.8 | - | - | 172.2 | / |
| Asparagine | N | 52.5 | 39.5 | / | / | / | 174.0 | 178.7 |
| Glycine | G | 43.6 | / | / | / | / | 171.3 | / |
| Phenylalanine | $F^{e}$ | 57.4 | 37.2 | 136.4 | - | - | 173.5 | / |
| Proline | $P^{a}$ | 61.4 | 30.1 | 25.5 | 48.9 | / | 175.0 | / |
| | $P^{b}$ | 60.5 | 31.0 | 25.9 | 48.7 | / | 173.9/175.1 | / |
| Tryptophan | $W^{a}$ | 55.0 | 28.4 | 130.6 | - | - | 175.0 | / |
| | $W^{b}$ | 55.7 | 28.4 | 130.7 | - | - | 175.8 | / |
| | $W^{c}$ | 54.5 | 28.5 | 129.9 | - | - | 176.5 | / |
| Valine | V | 56.4 | 29.2 | 19.2 | / | / | 178.7 | / |

**Supplementary Table S6. Molar composition of polysaccharides in cell walls.** The molar percentages of rigid cell-wall polysaccharides are estimated using integrals (volume) of cross peaks in 2D $^{13}C$-$^{13}C$ 53-ms CORD. The molar percentages of mobile cell-wall polysaccharides are estimated by using well-resolved peaks (C1 and C2) of 2D J-based $^{13}C$-$^{13}C$ DP-INADEQUATE. All errors are propagated from at least 3 individual integrals of well-resolved peaks for each type of molecule.

| Rigid domain | | | |
|---|---|---|---|
| Polysaccharide | Chitin% | Chitosan% | β-1,3-glucan |
| *R. delemar*, apo | 50 ± 3 | 45 ± 7 | 5 ± 1 |
| *R. delemar*, + nikko | 66 ± 3 | 34 ± 3 | / |
| *R. microsporus* | 76 ± 6 | 24 ± 5 | / |
| *M. lusitanicus* | 73 ± 5 | 27 ± 4 | / |
| *M. circinelloides* | 65 ± 8 | 35 ± 9 | / |
| *M. hiemalis* | 68 ± 7 | 32 ± 13 | / |

| Mobile domain | | | | |
|---|---|---|---|---|
| | *R. delemar*, apo | | *R. delemar*, +nikko | |
| Polysacharide | % | Peak | % | Peak |
| Chitin | 14.3 | C1, C2 | 2.5 | C1, C2 |
| Chitosan | 11.9 | C1, C2 | 9.4 | C1, C2 |
| N-acetylgalactosamine | 8.4 | C1, C2 | 0.9 | C1, C2 |
| Galactosamine | 7.5 | C1, C2 | 11.7 | C1, C2 |
| Galactopyranose | 19.2 | C1, C2 | 20.8 | C1, C2 |
| Galfuranose | 0.8 | C1, C2 | 0.5 | C1 |
| α-1,6-mannose | 4.7 | C1, C2 | 3.8 | C1, C2 |
| α-1,2-mannose | 15.3 | C1, C2 | 14.8 | C1, C2 |
| β-1,3-glucan | 4.7 | C1, C2 | 5.1 | C1, C2 |
| α-1,3-fucose | 10.4 | C1, C2 | 16 | C1, C2 |
| Unknown | 2.8 | C3, C4 and C5 | 14.6 | C3, C4, C5, C1 and C2 |

The following peaks were used for the estimation of molar composition of each rigid polysaccharide in fungal cell walls: ***R. delemar***: $Ch^{a,b}$-C4-1, C1-4, C4-5+C4-3, C5-4+C3-4; $Ch^{c}$-C1-6, C6-1, C6-2 and C2-6; $Ch^{d}$-ChCO-1, ChCO-4, ChCO-5, ChCO-3, ChCO-2, ChCO-6; $Cs^{a}$-C1-5, C1-3, C1-2 and C4-3; $Cs^{b}$-C1-5, C1-3, C1-2, C3-6 and C4-6; $Cs^{c}$-C1-5, C1-3, C1-2 and C1-4; $Cs^{d}$-C1-5, C1-3, C1-2 and C1-6; B-C1-4, C1-5, C5-4, C2-4, C3-5, C3-2 and C3-4. ***R. delemar* + nikko**: $Ch^{a,b}$-C4-1, C2-4, C6-2 and C5-4; $Ch^{d}$-ChCO-1, ChCO-4, ChCO-5 and ChCO-6; $Cs^{a}$-C3-4, C5-4, C4-6 and C4-3; $Cs^{b}$-C3-4, C4-6, C5-4 and C1-2; $Cs^{d}$-C4-1, C5-1, C3-1 and C2-1. ***R. microspores***: $Ch^{a,b}$-C4-1, C1-4, C1-5, and C5-4; $Cs^{a}$-C3-1, C4-1, C3-4, C5-4, and C4-5; $Cs^{b}$-C2-1, C1-2, C3-5 and C4-5. ***M. lusitanicus***: $Ch^{a,b}$-C4-1, C1-4, C5-4 and C4-5; $Cs^{a}$-C3-4, C5-4, C4-5 and C3-1; $Cs^{f}$-C4-2, C3-4, C2-1 and C1-2. ***M. cirinelloides***: $Ch^{a,b}$-C4-1, C1-4, C2-1 and C6-5; $Cs^{a}$-C3-4, C5-4, C4-5 and C4-3; $Cs^{f}$-C3-4, C4-3, C4-5 and C2-1. ***M. hiemalis***: $Ch^{a,b}$-C4-1, C1-4, C5-1, C4-6 and C4-2; $Ch^{d}$-ChCO-1, ChCO-4, ChCO-3 and ChCO-2. $Cs^{a}$-C2-1, C1-2, C5-4 and C3-4; $Cs^{e}$-C2-1, C1-2, C3-4 and C5-4.

**Supplementary Table S7. Intermolecular cross peaks in apo and nikkomycin-treated *R. delemar*.** The chemical shifts for the two dimensions of the spectra ($\omega_1$ and $\omega_2$). Asterisks indicate interactions where only one peak is observed along the diagonal line with the absence of the other.

| ***R. delemar*, apo** | | |
|---|---|---|
| Interaction | $\omega_1$ or $\omega_2$ (ppm) | $\omega_1$ or $\omega_2$ (ppm) |
| Chitin-chitosan | | |
| Ch$^{a,b}$Me-Cs$^{a}$1 | 22.9 | 101.9 |
| Ch$^{a,b}$Me-Cs$^{b}$1 | 22.9 | 98.1 |
| Ch$^{a,b}$Me-Cs$^{d}$1 | 22.9 | 100.7 |
| Ch$^{a,b}$Me-Cs4 | 22.9 | 79.7 |
| Cs$^{a}$3-Ch$^{a,b}$Me* | 72.6 | 22.9 |
| Cs$^{b}$3-Ch$^{a,b}$Me* | 71.1 | 22.9 |
| Ch$^{a,b}$Me-Cs3* | 22.9 | 70.7 |
| Ch$^{a,b}$Me-Cs$^{b}$2 | 22.9 | 56.5 |
| Ch2-Cs$^{b}$1 | 55.3 | 98.1 |
| Ch$^{c}$6-Cs$^{b}$1 | 58.9 | 98.1 |
| Ch$^{a,b}$6-Cs$^{a}$1 | 60.5 | 101.9 |
| Ch$^{a,b}$6-Cs$^{b}$1 | 60.5 | 98.1 |
| Ch$^{a,b}$6-Cs$^{d}$1 | 60.5 | 100.7 |
| Ch$^{c}$6-Cs4* | 58.9 | 79.5 |
| Ch2-Cs4 | 55.3 | 79.5 |
| Cs3-Ch$^{a,c}$4 | 70.6 | 83.3 |
| Cs3-Ch$^{b}$4 | 70.6 | 84.3 |
| Ch3-Cs$^{b}$1 | 74.1 | 98.1 |
| Ch3-Cs$^{a}$1 | 74.1 | 101.9 |
| Ch$^{a,c}$4-Cs$^{a}$1 | 83.3 | 101.9 |
| Ch$^{b}$4-Cs$^{a}$1 | 84.3 | 101.9 |
| Ch1-Cs4 | 104.2 | 79.5 |
| Ch$^{a,c}$4-Cs$^{d}$1 | 83.3 | 100.7 |
| Ch$^{b}$4-Cs$^{d}$1 | 84.3 | 100.7 |
| Cs$^{b}$1-Ch1 | 98.1 | 104.2 |
| Cs$^{a}$1-Ch1 | 101.9 | 104.2 |
| Cs$^{d}$1-Ch1 | 100.7 | 104.2 |
| Cs$^{b}$3-ChCO | 71.1 | 174.9 |
| Cs$^{a}$3-ChCO | 72.6 | 174.9 |
| Cs$^{a}$1-ChCO | 101.9 | 174.9 |
| Cs$^{b}$1-ChCO | 98.1 | 174.9 |
| Chitin-Chitin | | |
| Ch$^{a}$6-Ch$^{b}$4 | 60.5 | 84.3 |
| Ch$^{a,b}$Me-Ch$^{c}$6 | 22.9 | 60.5 |
| Ch$^{c}$6-Ch$^{a,b}$6 | 58.9 | 60.5 |
| Chitosan-Chitosan | | |
| Cs$^{b}$1-Cs$^{a}$1 | 98.1 | 101.9 |
| Cs$^{b}$6-Cs$^{a}$1 | 60.7 | 101.9 |
| Chitin-β-1,3-glucan | | |
| Ch$^{a,b}$Me-B5 | 22.9 | 77.4 |
| Ch$^{c}$6-B5 | 58.9 | 77.4 |
| B5-ChCO | 77.4 | 174.9 |
| B4-Ch$^{a,c}$5 | 68.2 | 75.7 |
| Chitosan-β-1,3-glucan | | |
| B5-Cs$^{a}$1 | 77.4 | 101.9 |
| B5-Cs$^{b}$1 | 77.4 | 98.1 |
| B5-Cs$^{d}$1 | 77.4 | 100.7 |
| B4-Cs$^{a}$1* | 68.2 | 101.9 |
| B4-Cs$^{b}$1* | 68.2 | 98.1 |
| B4-Cs4* | 68.2 | 79.5 |
| Carbohydrates-Protein | | |
| Iγ2-Cs$^{a}$1* | 15.5 | 101.9 |
| Iγ2-Cs$^{b}$1* | 15.5 | 98.4 |
| Iγ2-Cs4* | 15.5 | 79.5 |
| Iγ2-Cs$^{a}$5$^{*}$ | 15.5 | 75.0 |
| Iγ2-Ch3* | 15.5 | 74.0 |
| Iγ2-Ch5* | 15.5 | 75.1 |
| Iγ1-Ch1 | 25.6 | 104.2 |
| Iγ1-Ch3 | 25.6 | 74.0 |

| ***R. delemar*, + nikko** | | |
|---|---|---|
| Interaction | $\omega_1$ or $\omega_2$ (ppm) | $\omega_1$ or $\omega_2$ (ppm) |
| Chitin-chitosan | | |
| ChMe-Cs$^{a}$1 | 22.8 | 102.2 |
| ChMe-Cs$^{d}$1 | 22.8 | 101.6 |
| ChMe-Cs$^{b}$1 | 22.8 | 99.8 |
| ChMe-Cs$^{a,b}$2 | 22.8 | 57 |
| ChMe-Cs$^{a,b}$3 | 22.8 | 72.4 |
| ChMe-Cs$^{a}$4 | 22.8 | 79.9 |
| ChMe-Cs$^{b}$4 | 22.8 | 77.6 |
| ChMe-Cs$^{a,b}$6 | 22.8 | 60.8 |
| Ch1-Cs$^{a}$1 | 104.4 | 102.2 |
| Ch1-Cs$^{d}$1 | 104.4 | 101.6 |
| Ch1-Cs$^{b}$1 | 104.4 | 99.8 |
| Ch1-Cs$^{b}$2 | 104.4 | 57.2 |
| Ch1-Cs$^{b}$3 | 104.4 | 72.4 |
| Ch2-Cs$^{a}$1 | 55.2 | 102.2 |
| Ch2-Cs$^{b}$1 | 55.2 | 57.0 |
| Ch3-Cs$^{b}$1 | 83.2 | 99.8 |
| Ch4-Cs$^{a}$1 | 83.2 | 102.2 |
| Ch4-Cs$^{b}$1 | 83.2 | 99.8 |
| Ch4-Cs$^{a,b}$5 | 83.2 | 75.3 |
| Ch4-Cs$^{a}$3* | 83.2 | 72.8 |
| Ch4-Cs$^{b}$3* | 83.2 | 72.4 |
| Cs$^{a,b}$3-Ch4* | 72.4 | 83.2 |
| Ch5-Cs$^{b}$1 | 76.3 | 102.2 |
| Ch6-Cs$^{b}$1 | 60.5 | 99.8 |
| ChCO-Cs$^{a}$1 | 174.8 | 102.2 |
| ChCO-Cs$^{d}$1 | 174.8 | 101.6 |
| ChCO-Cs$^{b}$1 | 174.8 | 99.8 |
| ChCO-Cs$^{a,b}$3 | 174.8 | 72.4 |
| ChCO-Cs$^{a}$4 | 174.8 | 79.9 |
| ChCO-Cs$^{b}$4 | 174.8 | 77.6 |
| Ch4-Cs$^{b}$2 | 83.2 | 57.0 |
| Ch1-Cs$^{a}$4 | 104.4 | 79.9 |
| Ch5-Cs$^{a}$1 | 76.3 | 102.2 |
| Chitosan-chitosan | | |
| Cs$^{a}$2-Cs$^{b}$1 | 56.1 | 99.8 |
| Cs$^{a}$4-Cs$^{b}$1 | 79.9 | 99.8 |
| Cs$^{a}$4-Cs$^{b}$3 | 79.9 | 72.4 |
| Cs$^{a}$5-Cs$^{b}$1 | 75.3 | 99.8 |
| Cs$^{a}$1-Cs$^{b}$1 | 102.2 | 99.8 |
| Cs$^{a}$1-Cs$^{b}$2 | 102.2 | 57 |
| Carbohydrates-Protein | | |
| Iγ1-Ch1 | 25.8 | 104.4 |
| Iγ1-Ch4 | 25.8 | 83.3 |
| Iγ1-Ch5 | 25.8 | 76.4 |
| Iγ1-Ch3 | 25.8 | 73.7 |
| Iγ1-Ch6* | 25.8 | 61.0 |
| Cs$^{b}$3-Iγ2* | 15.9 | 72.4 |
| Cs$^{b}$2-Iγ2* | 15.9 | 57.0 |

**Supplementary Table S8. Water-edited intensities of polysaccharide carbon sites.** The intensity ratios were calculated by comparing the peak intensity in water-edited and control spectra, which were normalized by the number of scans. The error bars represent the standard deviations propagated from NMR signal-to-noise ratios.

| Cross peaks | apo | + nikko | Cross peaks | apo | Cross peaks | + Nikko | Cross peaks | apo | Cross peaks | + Nikko |
|---|---|---|---|---|---|---|---|---|---|---|
| **Average Ch[a,b]** | **0.24** | **0.36** | **Average Cs[a]** | **0.59** | **Average Cs[a]** | **0.50** | **Average Cs[b]** | **0.47** | **Average Cs[b]** | **0.49** |
| Ch1-1 | 0.17±0.04 | 0.35±0.01 | Cs1-1 | 0.86±0.06 | Cs1-1 | 0.42±0.04 | Cs1-1 | 0.58±0.10 | Cs1-1 | 0.39±0.02 |
| Ch1-4 | 0.15±0.04 | 0.33±0.03 | Cs1-2 | 0.38±0.05 | Cs1-2 | 0.52±0.06 | Cs1-2 | 0.28±0.05 | Cs1-4 | 0.58±0.05 |
| Ch1-5 | 0.17±0.03 | 0.37±0.03 | Cs1-3 | 0.55±0.05 | Cs1-3 | 0.57±0.05 | Cs1-3 | 0.62±0.06 | Cs1-5 | 0.55±0.07 |
| Ch1-3 | 0.18±0.03 | 0.37±0.02 | Cs1-4 | 0.78±0.06 | Cs1-4 | 0.61±0.07 | Cs1-4 | 0.56±0.03 | Cs1-3 | 0.63±0.06 |
| Ch1-6 | 0.20±0.05 | 0.43±0.07 | Cs1-5 | 0.78±0.05 | Cs1-5 | 0.60±0.05 | Cs1-5 | - | Cs1-2 | 0.48±0.06 |
| Ch1-2 | 0.17±0.02 | 0.34±0.02 | Cs1-6 | 0.44±0.02 | Cs1-6 | - | Cs1-6 | 0.45±0.10 | Cs4-1 | 0.55±0.08 |
| Ch4-1 | 0.14±0.03 | 0.34±0.05 | Cs5-1 | 0.55±0.04 | Cs3-1 | 0.58±0.09 | Cs4-1 | 0.35±0.06 | Cs4-4 | 0.46±0.03 |
| Ch4-4 | 0.28±0.04 | 0.34±0.03 | Cs5-2 | 0.54±0.03 | Cs3-4 | 0.43±0.05 | Cs4-2 | 0.39±0.04 | Cs4-5 | 0.40±0.08 |
| Ch4-5 | 0.21±0.03 | 0.31±0.04 | Cs5-3 | 0.39±0.06 | Cs3-5 | 0.49±0.04 | Cs4-3 | 0.36±0.02 | Cs4-3 | 0.38±0.07 |
| Ch4-3 | 0.25±0.07 | 0.33±0.04 | Cs5-4 | 0.66±0.03 | Cs3-3 | 0.51±0.07 | Cs4-4 | 0.59±0.05 | Cs4-2 | 0.46±0.13 |
| Ch4-6 | 0.20±0.02 | 0.32±0.07 | Cs5-5 | 0.41±0.01 | Cs3-2 | - | Cs4-5 | 0.48±0.06 | - | - |
| Ch4-2 | 0.30±0.04 | 0.36±0.05 | Cs5-6 | 0.45±0.03 | Cs2-1 | 0.54±0.10 | Cs4-6 | 0.52±0.08 | Cs5-1 | 0.45±0.11 |
| Ch2-1 | 0.18±0.02 | 0.31±0.02 | Cs6-1 | 0.87±0.04 | Cs2-4 | 0.45±0.09 | Cs5-1 | 0.40±0.03 | Cs5-4 | 0.45±0.08 |
| Ch2-4 | 0.29±0.03 | 0.35±0.04 | Cs6-2 | 0.61±0.05 | Cs2-5 | 0.47±0.06 | Cs5-2 | 0.44±0.03 | Cs5-5 | 0.45±0.03 |
| Ch2-5 | 0.28±0.02 | 0.37±0.02 | Cs6-3 | 0.40±0.04 | Cs2-3 | 0.45±0.06 | Cs5-3 | 0.37±0.01 | Cs5-3 | 0.42±0.15 |
| Ch2-3 | 0.21±0.02 | 0.33±0.01 | Cs6-4 | 0.81±0.10 | Cs2-2 | 0.45±0.04 | Cs5-4 | 0.65±0.04 | Cs5-2 | 0.46±0.10 |
| Ch2-6 | 0.21±0.03 | 0.34±0.04 | Cs6-5 | 0.62±0.08 | - | - | Cs5-5 | 0.24±0.01 | - | - |
| Ch2-2 | 0.27±0.03 | 0.37±0.02 | Cs6-6 | 0.60±0.07 | - | - | Cs5-6 | 0.43±0.04 | Cs2-1 | 0.46±0.07 |
| Ch3-1 | 0.20±0.03 | 0.31±0.02 | Cs4-1 | - | Cs4-1 | 0.44±0.06 | Cs2-1 | 0.39±0.03 | Cs2-4 | 0.46±0.08 |
| Ch3-4 | 0.27±0.06 | 0.35±0.02 | Cs4-2 | - | Cs4-2 | 0.55±0.08 | Cs2-2 | 0.59±0.09 | Cs2-5 | 0.81±0.13 |
| Ch3-5 | 0.41±0.03 | 0.43±0.01 | Cs4-3 | - | Cs4-3 | 0.43±0.04 | Cs2-3 | 0.52±0.09 | Cs2-3 | 0.49±0.13 |
| Ch3-3 | 0.50±0.04 | 0.42±0.01 | Cs4-4 | - | Cs4-4 | 0.52±0.03 | Cs2-4 | 0.65±0.06 | Cs2-2 | 0.47±0.05 |
| Ch3-6 | 0.37±0.03 | 0.54±0.03 | Cs4-5 | - | Cs4-5 | 0.48±0.05 | Cs2-5 | 0.47±0.06 | - | - |
| Ch3-2 | 0.18±0.02 | 0.37±0.02 | Cs4-6 | - | Cs4-6 | - | Cs2-6 | 0.48±0.03 | - | - |

| Cross peaks | apo | Cross peaks | apo | Cross peaks | apo |
|---|---|---|---|---|---|
| **Average Ch[c]** | **0.27** | **Average Cs[c]** | **0.71** | **Average β-1,3-glucan** | **0.29** |
| Ch1-Ch1 | 0.17±0.04 | Cs1-1 | 0.83±0.07 | B3-1 | 0.31±0.04 |
| Ch1-Ch4 | 0.24±0.03 | Cs1-2 | 0.66±0.12 | B3-2 | 0.13±0.04 |
| Ch1-Ch3 | 0.22±0.03 | Cs1-3 | 0.89±0.17 | B3-3 | 0.28±0.04 |
| Ch1-Ch6 | 0.50±0.07 | Cs1-4 | 0.84±0.08 | B3-4 | 0.16±0.03 |
| Ch1-Ch2 | 0.18±0.06 | Cs1-5 | 0.56±0.08 | B3-5 | 0.12±0.07 |
| Ch4-Ch1 | 0.34±0.06 | Cs5-1 | 0.62±0.07 | B5-1 | 0.24±0.09 |
| Ch4-Ch4 | 0.28±0.03 | Cs5-2 | 0.64±0.05 | B5-2 | 0.33±0.04 |
| Ch4-Ch3 | 0.27±0.05 | Cs5-3 | 0.38±0.05 | B5-3 | 0.41±0.08 |
| Ch4-Ch6 | 0.33±0.11 | Cs5-4 | 0.74±0.09 | B5-4 | 0.45±0.06 |
| Ch4-Ch2 | 0.25±0.07 | Cs5-5 | 0.39±0.03 | B5-5 | 0.44±0.08 |
| Ch2-Ch1 | 0.18±0.02 | Cs3-1 | 0.91±0.05 | B4-1 | 0.14±0.02 |
| Ch2-Ch4 | 0.26±0.07 | Cs3-2 | 0.69±0.05 | B4-2 | 0.22±0.07 |
| Ch2-Ch3 | 0.30±0.06 | Cs3-3 | 0.88±0.06 | B4-3 | 0.31±0.02 |
| Ch2-Ch6 | 0.24±0.07 | Cs3-4 | 0.89±0.08 | B4-4 | 0.35±0.04 |
| - | - | Cs3-5 | 0.73±0.04 | B4-5 | 0.40±0.04 |

**Supplementary Table S9. $^{13}$C-$T_1$ and $^{1}$H-$T_{1\rho}$ relaxation time constants and dipolar order parameters.** Single exponential equations were used to fit the observed de cay: $I(t) = e^{-t/T_1}$. The fit parameters were obtained, and the standard deviations of these parameters were calculated as error bars. Average values are shown in bold.

| ***R. delemar*, apo** | | | | |
|---|---|---|---|---|
| Carbohydrate | | Chemical shifts | $^{13}$C-$T_1$ Time (s) | $^{1}$H-$T_{1\rho}$ Time (ms) |
| chitin | | **Average** | **1.7** | **5.6** |
| | a,b | 104 | 2.4±0.1 | 8.8±0.4 |
| | b | 84.0 | 2.3±0.1 | 7.3±0.6 |
| | a,c | 83.3 | 2.2±0.2 | 7.8±0.5 |
| | a,c | 75 | 1.1±0.1 | 2.7±0.2 |
| | a | 60.5 | 0.9±0.1 | 3.0±0.2 |
| | c | 59 | 1.3±0.1 | 3.4±0.2 |
| | a,b,c | 55.5 | 1.5±0.1 | 5.7±0.5 |
| chitosan | | **Average** | **0.74** | **1.9** |
| | a | 102 | 0.95±0.09 | 1.4±0.1 |
| | b | 98.1 | 0.84±0.06 | 1.3±0.1 |
| | a,b | 79.5 | 0.63±0.03 | 1.5±0.1 |
| | a | 72.6 | 0.61±0.05 | 3.1±0.3 |
| | b | 71.2 | 0.68±0.05 | 2.0±0.1 |
| β-1,3-glucan | | **Average** | **0.65** | **1.8** |
| | | 77.5 | 0.51±0.03 | 1.3±0.1 |
| | | 68.1 | 0.67±0.1 | 1.6±0.1 |
| | | 61.3 | 0.78±0.09 | 2.6±0.2 |

| ***R. delemar*, nikko** | | | | |
|---|---|---|---|---|
| chitin | | **Average** | **2.6** | **12** |
| | a,b | 104.2 | 4.0±0.2 | 15±2 |
| | a,b | 83.3 | 4.0±0.2 | 11±2 |
| | a,b | 73.6 | 2.2±0.3 | 6.9±0.9 |
| | a,b | 60.9 | 1.0±0.2 | 4.1±0.3 |
| | a,b | 55.4 | 2.0±0.2 | 5.5±0.5 |
| chitosan | | **Average** | **1.1** | **5.3** |
| | a | 102.2 | 1.4±0.09 | 4.3±0.2 |
| | a | 79.9 | 0.93±0.07 | 3.8±0.3 |
| | a | 72.8 | 1.6±0.2 | 8.3±0.5 |
| | a | 75.3 | 1.7±0.2 | 8.3±0.5 |
| | b | 99.8 | 0.80±0.04 | 3.2±0.2 |
| | b | 77.6 | 0.84±0.06 | 4.1±0.2 |
| | b | 72.4 | 0.87±0.11 | 4.7±0.4 |
| | b | 57.0 | 0.99±0.08 | 4.4±0.2 |
